# Tag with Caution — How protein tagging influences the formation of condensates

**DOI:** 10.1101/2024.10.04.616694

**Authors:** Kerstin Dörner, Michelle Gut, Daan Overwijn, Fan Cao, Matej Siketanc, Stephanie Heinrich, Nicole Beuret, Timothy Sharpe, Kresten Lindorff-Larsen, Hondele Maria

## Abstract

Fluorescent proteins and peptide tags are essential tools in cellular biology, but can alter the biochemical properties of target proteins. Biomolecular condensates, which have emerged as key principles of cellular organization, are suggested to provide robustness to cells, yet they can also respond sensitively to small changes in environmental conditions—or tagging of their components, as our findings suggest. Here, we investigated the effects of sixteen widely used tags on condensate formation in various model organisms, *in vitro*, in cells and by computational modelling. We find that tagging strongly influenced condensation for some proteins, while others remained unaffected. Effects varied, with some tags enhancing and others decreasing condensation, and depended on the protein being tagged. Coarse-grained simulations suggest that the charge of the fluorescent protein tags is a critical factor modulating condensation behavior. Together, our results underscore the importance of rigorous experimental design and interpretation in condensate experiments.

## Introduction

Genetically encoded protein and peptide tags have become essential tools in studying protein localization and function, providing an indispensable toolkit for modern cell biology. Widely used for applications such as affinity purification, microscopy, proximity labeling, and targeted protein degradation^1^, these tags offer numerous advantages. However, they also pose risks. Tagging introduces steric hindrance and altered or additional chemical and surface properties, which can perturb a protein’s physiological behavior, affecting its interactions, folding, turnover, enzymatic activity, and localization^2^.

A study comparing the localization of roughly 500 human proteins using either immunostaining or fluorescent protein (FP) tagging revealed that approximately 20% of the proteins exhibit different localization patterns^3^. Also, immunostaining has to be performed with care: fixation and permeabilization processes can affect the perceived distribution of proteins^4^, and individual antibodies may lack specificity.

FP tags are widely used in the study of biomolecular condensates both *in vitro* and in cells, such as nucleoli, nuclear speckles (NS), P-bodies (PB) or stress granules (SG). Condensate formation often relies on transient, relatively weak, multivalent interactions, typically resulting in dynamic and viscoelastic structures with rapid movement of components within the condensate and exchange between the condensate and its surroundings. Condensation is influenced by various factors, including protein concentration, posttranslational modifications, and environmental conditions such as pH or salt concentration^5–8^. However, the inherently transient and weak nature of these interactions also makes condensates particularly vulnerable to experimental artifacts and perturbations.

Despite their widespread use, the impact of FP and peptide tags on protein condensation remains somewhat anecdotal and is still not well-documented. Recent studies have begun to shed light on how FP and other tags can considerably alter the condensation or aggregation behavior of specific proteins of interest (POI)^9,10^. For example, tagging huntingtin Httex1(Q25) with RFP promotes condensation in conditions where the untagged protein remains dispersed^11^. Similarly, mEGFP, FusionRed, mNeonGreen or Halo increase condensate size and protein density for viral µNS^12^, while AID-sfGFP or mEGFP increase and decrease condensation of HP1α respectively^13^. Quantitative *in vitro* measurements of phase separation showed differences in the salt-dependency of the propensity to phase separate between hnRNPA1 and a His-SUMO tagged fusion protein^14^, and a series of genetically encoded sensors for phase separation gave different responses depending on which FP was used^15^. These studies highlight that protein tags can introduce artifacts, yet a comprehensive comparative analysis of how different FP and other tags affect condensation is lacking.

We therefore systematically tested the influence of FP and peptide tags on several prominent condensation-prone proteins involved in cytoplasmic and nuclear condensate formation across human, *Saccharomyces cerevisiae*, and *Escherichia coli*, in cells, *in vitro* and using computational simulations. Initially, we examined 16 genetically encoded tags fused to DDX3X in HeLa cells, revealing changes in DDX3X SG appearance that correlated with altered condensation behavior of selected DDX3X-FP constructs *in vitro*. Further analysis of human condensate markers showed that while EDC3 (PB) was affected by FP tagging as well, G3BP1 (SG) and SRRM2 (NS) were not. Additionally, FP tagging influenced the condensation of *S. cerevisiae* Pab1, both *in vivo* and *in vitro*, and of *E. coli* CsdA and human FUS *in vitro*. Our experiments show that mCherry consistently reduces condensate formation, while EGFP or EYFP tend to induce larger and more aggregated structures. We also found that affinity tags commonly used for protein purification can alter RNA specificity in condensate formation. Our findings indicate that the tag effect is POI-dependent, and that some POI are more affected than others. These observations are also seen in molecular simulations that suggest that charge effects can explain some, but not all effects of tagging on the propensity to phase separate. This leads us to conclude that tagging should be avoided whenever possible or cautiously evaluated when experimentally required.

## Results

### Fluorescent protein, peptide or ligand-binding tags strikingly alter DDX3X stress granule appearance

To systematically investigate how tagging condensate marker proteins with various FP and peptide tags affects their appearance in cells, we first focused on DDX3X, a DEAD-box ATPase and key SG marker^16^. SG are cytoplasmic condensates (0.1-2 µm) that form in response to stressors like sodium arsenite or heat, and are enriched in mRNAs and RNA-binding proteins, including stalled translation pre-initiation complexes^17,18^. DDX3X itself forms condensates *in vitro*, and deletion of its N-terminal intrinsically disordered region (IDR) significantly reduces both condensation and SG formation in cells^19–22^.

Initially, we screened the effect of 16 tags C-terminally fused to DDX3X (Supplementary Table 1, Figure S1A-C): nine commonly used FP (ECFP, EGFP, mEGFP, mNeonGreen, EYFP, mYPet, mCherry, mScarlet-I, mRuby3), five peptide tags (myc, V5, HA, FLAG, and alfa), and two ligand-binding tags (Halo and SNAP) (Figure 1A). We transiently expressed these constructs in HeLa K cells, induced SG formation using sodium arsenite and fixed the cells 24 h post transfection. As a reference, we included an antibody staining against endogenous DDX3X. Compared to immunostained endogenous DDX3X, cells expressing tagged DDX3X constructs had considerably altered SG characteristics (Figure 1A/B). We quantified the change by measuring the number of SG foci per cell, the average area of individual foci, and an ‘intensity enrichment coefficient’ (IEC), calculated as the median intensity of each focus divided by the median intensity of the entire cell. Different FP induced strikingly different SG morphologies. ECFP, EGFP, EYFP, and mNeonGreen resulted in larger SG. For ECFP and EYFP the granules became non-spherical, potentially indicating protein aggregation. EGFP, EYFP and mNeonGreen increased SG numbers, suggesting that they enhance the accumulation of DDX3X fusion proteins in SG, which is reflected by higher IEC values compared to other FPs. However, for all FP the IEC was substantially decreased compared to the AB-stained endogenous DDX3X. mCherry and mScarlet-I reduced SG numbers while maintaining similar SG size, and mRuby3 showed SG numbers comparable to endogenous DDX3X but with slightly enlarged sizes. All three FP displayed very low IEC values, indicating impaired recruitment of DDX3X to SG (Figure 1B). In summary, FP tagging in most cases strongly altered the appearance of DDX3X SG. The best option in our opinion is mEGFP, which produced SG similar in size to endogenous DDX3X with only a slight increase in SG numbers and average IEC.

**Figure 1:**
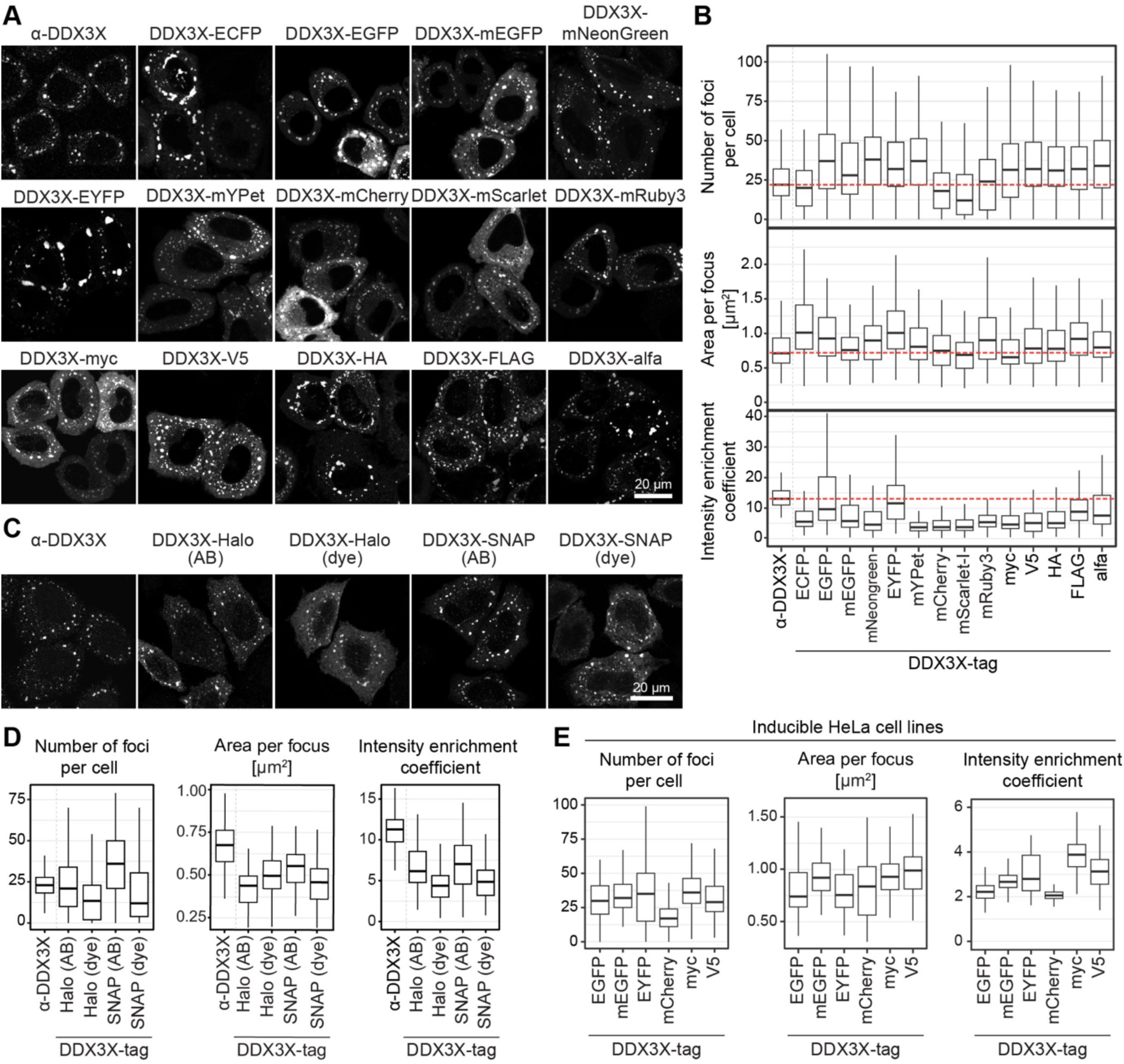
Expression of DDX3X with various tags influences stress granule formation in human cells Transient or stable inducible expression of DDX3X constructs for 24 h in HeLa cells. Cells were stressed with 500 μM sodium arsenite for 30 min before fixation. For each experiment three independent replicates were performed (N = 3). Quantification of DDX3X foci includes number of foci per cell, area per focus and intensity enrichment coefficient (IEC, median intensity per focus/ median intensity of the cell). (A) Transient expression of DDX3X fused with FP and peptide tags in HeLa K cells. Untransfected cells were immunostained for DDX3X. Peptide tags were visualized by immunostaining with the respective antibodies. (B) Quantification of (A); N = 3, n ≥ 135 cells. (C) Transient expression of DDX3X fused with Halo and SNAP tag in HeLa K cells. Tags were either visualized by incubation with respective dyes (dye) or by immunostaining (AB). (D) Quantification of (C); N = 3, n ≥ 90 cells. (E) Quantification of DDX3X foci in stable HeLa cell lines, induced with doxycycline to express DDX3X fused with FP and peptide at endogenous level as displayed in S1D/E. N = 3, n ≥ 65 cells.

All tested peptide tags increased the number of SG foci per cell (Figure 1A/B). Tagging with HA, FLAG, and alfa led to the formation of a few larger and irregularly shaped foci, indicating protein aggregation. Myc and V5 behaved quite similarly to the endogenous protein across all parameters, indicating they could be good choices for tagging in cells. Overall, peptide tags had a more consistent impact on DDX3X fusion constructs compared to FP tags.

Furthermore, we tested DDX3X tagged with the ligand-binding proteins Halo and SNAP, visualizing them using either anti-tag antibodies or tag-binding dyes (Figure 1C). The SG foci of DDX3X-Halo/SNAP were smaller and more irregular compared to endogenous DDX3X (Figure 1C/D), suggesting aggregation. Notably, visualization with dyes significantly reduced both SG numbers and IEC values, indicating impaired recruitment of DDX3X to the granules (Figure 1D). In conclusion, SNAP and Halo tags appear suboptimal for studying DDX3X in cells.

Given the variability in expression levels between cells from transient transfections, we next assessed how SG characteristics correlate with DDX3X expression levels (Figure S1D-F). While endogenous DDX3X displayed a relatively uniform distribution of SG number and area, transiently expressed tagged DDX3X showed greater variability of these parameters for each expression level bin. Surprisingly, for most tags, we observed only a weak correlation between expression levels and SG number or area.

### Titrating expression levels results in milder and less variable effects on DDX3X stress granule appearance

Considering the high expression and phenotypic variability observed in transiently transfected cells, we generated stable HeLa cell lines expressing DDX3X tagged with EGFP, mEGFP, EYFP, mCherry, myc, or V5 from a doxycycline-inducible promoter. We titrated expression levels to match endogenous DDX3X at a 1:1 ratio (Figure S1G/H). These cells largely mirrored the trends from transient transfections, though the effects were considerably milder (Figure 1E). mEGFP, EGFP, EYFP, myc and V5 showed slightly increased foci numbers compared to endogenous DDX3X, with mEGFP also showing a somewhat larger area and myc and V5 increased IEC values (Figure 1E). Strikingly, DDX3X-mCherry showed a drastic reduction of SG numbers even at endogenous expression levels. In this experiment, we additionally immunostained for another SG marker, G3BP1, which showed minimal differences in foci between cells expressing DDX3X with no or different FP tags (Figure S1G).

In summary, DDX3X-mCherry affects SG also at endogenous expression levels, whereas the effects of other tags can become negligible. This suggests that even low levels of transient expression may still represent significant overexpression, contributing to the observed phenotypes.

### General considerations for studying condensation *in vitro*

We wondered whether the effects we observed in cells might originate from inherent differences in the ability of different DDX3X fusion proteins to form condensates. To investigate this, we purified DDX3X both untagged and tagged with various FP (Figure S2A). Condensates formed from untagged proteins can be visualized by brightfield or quantitative phase microscopy^23^. However, the use of widely accessible widefield or confocal microscopy presents several challenges. Critical biophysical parameters, such as the partition coefficient (PC), which reflects the relative protein concentration inside and outside the condensates, and fluorescence recovery after photobleaching (FRAP), used to assess component turnover dynamics, cannot be determined. Additionally, multi-component condensates cannot easily be analyzed, and untagged condensates present difficulties in maintaining proper focus during automated microscopy workflows. Therefore, we introduced a spike-in of 1% POI chemically labeled at the N-terminus with ATTO NHS-ester dyes. Given that we observed no differences in condensation behavior between the untagged and untagged+ATTO samples for DDX3X (Figure S2B), we designated untagged+ATTO as the reference point for all subsequent *in vitro* experiments. For calculations of PC and FRAP analysis we assumed that the untagged+ATTO samples would behave exactly like untagged samples.

Also, it is important to keep in mind that certain FP are pH-sensitive. For example, at lower pH levels (pH 6-7), EGFP and EYFP, both of which are moderately acid-sensitive, appear much dimmer compared to their appearance at higher pH levels (pH 7-8)^24^. In this manuscript we display images with identical exposure levels for all panels of one FP, except for Figure 6 where panels were so different that we adjusted exposure levels individually for each image.

As previously reported^25^, condensates undergo growth and fusion over time (Figure S2C). Therefore, within an experiment it is critical to precisely control and standardize the time between condensate formation and image acquisition for each individual sample to ensure consistency in experimental observations.

### Fluorescent protein tags modify the characteristics of *in vitro* DDX3X condensates

We examined the condensation behavior of DDX3X with poly-uridylic acid (poly(U)) across varying pH and salt conditions (Figure 2A-D). Untagged+ATTO DDX3X formed condensates at pH 6.2 but tended to aggregate at higher pH (Figure 2A). These condensates were highly sensitive to salt, visible at 100 mM NaCl, shrinking at 200 mM, and disappearing at 400 mM (Figure 2C/D, S2D/E).

**Figure 2:**
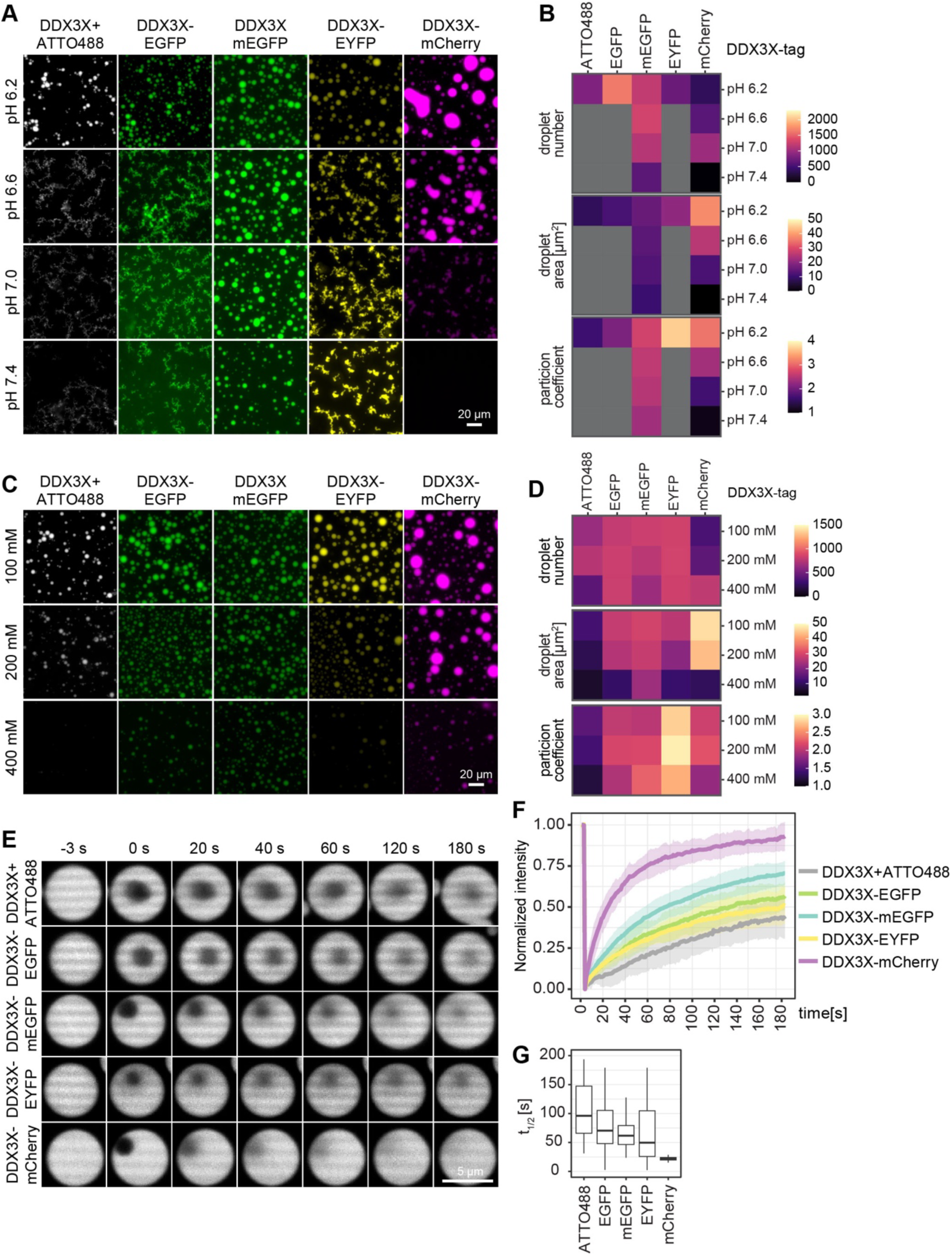
Fluorescent protein tags influence DDX3X condensate formation and turnover *in vitro* DDX3X *in vitro* condensation assays with untagged+1% ATTO488 spike-in, or FP-tagged proteins were incubated at 25°C for 30 min before imaging. Quantification includes number of condensates per 0.1 mm^2^, droplet area and mean PC. (A) 5 µM DDX3X in 25 mM sodium phosphate buffer at the indicated pHs, 150 mM NaCl, 2 mM MgCl_2_, 0.5 mg/mL BSA, 0.05 mg/mL poly(U), 0.5 mM ATP, 0.5 mM DTT. Incubated at 25°C for 30 min before imaging. (B) Quantification of (A): Images displaying aggregates were not quantified and are shown in grey. 4 well positions were imaged in three independent replicates. (C) 5 µM DDX3X in 25 mM sodium phosphate buffer at pH 6.2, at different salt concentrations (100, 200 or 400 mM NaCl), 2 mM MgCl_2_, 0.5 mg/mL BSA, 0.05 mg/mL poly(U), 0.5 mM ATP, 0.5 mM DTT. (D) Quantification of (C): 4 well positions were imaged in three independent replicates. (E) FRAP of *in vitro* DDX3X condensates of similar size. 5 µM DDX3X in 25 mM sodium phosphate buffer at the pH 6.2, 100 mM NaCl, 2 mM MgCl_2_, 0.5 mg/mL BSA, 0.05 mg/mL poly(U), 0.5 mM ATP, 0.5 mM DTT. Condensates were matured for 1 h at 25°C before FRAP. Fluorescence recovery curve of DDX3X droplets (**F**) and time till half recovery of DDX3X droplets (**G**) were analyzed. N = 3, n ≥ 23 condensates.

When DDX3X was tagged with the highly related EGFP or EYFP (Figure S1A-C), it showed similar behavior but typically formed larger condensates with higher PC and increased salt resistance. DDX3X tagged with mEGFP formed larger, round condensates across all pH conditions, showing less sensitivity to pH and salt compared to untagged+ATTO DDX3X and EGFP. The mCherry tag overall had the biggest influence on the appearance of DDX3X droplets. DDX3X-mCherry produced the largest and ‘roundest’ condensates at low salt and pH, suggesting enhanced fusion, but these shrank and disappeared at higher pH and salt levels (Figure 2C/D). In summary, these findings indicate that FP tags considerably affect the appearance of DDX3X condensates *in vitro*, suggesting that the SG phenotypes observed in cells may be at least partly driven by altered condensation behavior of DDX3X itself in addition to potential interactions between the FPs and other cellular components.

Since some FP are known to dimerize^24^, we conducted analytical ultracentrifugation (AUC) in the same buffers used for DDX3X condensation assays (150 mM NaCl, pH 7.4 and 6.2) (Figure S2F/G). In both conditions, EGFP and EYFP showed partial dimerization as previously reported, while mEGFP and mCherry remained monomeric. This dimerization likely contributes to the pronounced effects of EGFP and EYFP on DDX3X condensation, but is probably just one of several factors, including surface charge (Figure S2A/B).

To qualitatively evaluate the material properties of DDX3X-mCherry versus untagged+ATTO, we conducted a time-course analysis of condensate formation (Figure S2C). Both proteins showed condensate fusion, but DDX3X-mCherry formed larger condensates while smaller ones completely disappeared, whereas untagged+ATTO DDX3X retained smaller, more solid-like condensates.

To investigate potential differences in material properties in more detail, we analyzed condensate turnover using FRAP for untagged+ATTO and FP-tagged DDX3X (Figure 2E/F). All proteins showed recovery of the bleached area, however, recovery kinetics varied: untagged+ATTO DDX3X had a relatively slow recovery (t_1/2_ ∼ 100 s), while FP-tagged variants recovered faster: EGFP, mEGFP, and EYFP with t_1/2_ between 50-70 s, and mCherry with a t_1/2_ of ∼25 s (Figure 2G, S2H). Moreover, FP tags appeared to influence the mobile fraction of DDX3X, with mCherry-tagged DDX3X achieving full recovery within 180 s, while the other curves approached considerably lower plateau levels. This suggests that a portion of the proteins are less mobile and potentially aggregated. These findings align with the observation that untagged+ATTO, EGFP, and EYFP tagged DDX3X show aggregation at higher pH, and indicate that aggregated states may be present at pH 6.2 already, where FRAP was conducted.

In conclusion, FP tags have a substantial impact on DDX3X condensate appearance. At physiological pH, DDX3X-mCherry condensates do not form, both *in vitro* and in cells. In contrast, EYFP and EGFP tags promote aggregation and reduce FRAP recovery *in vitro*, which might explain the enlarged SG observed in cells upon transient (over)expression. This highlights the importance of screening FP tags *in vitro* when conducting cellular experiments to select those that minimally affect protein behavior.

### mCherry-EDC3 strongly reduces P-body numbers and DCP1 recruitment

We wondered whether the effects of FP tagging observed with DDX3X extend to other condensation-prone proteins. To explore this, we used a smaller set of four FP (EGFP, mEGFP, EYFP, mCherry) and two peptide tags (V5, myc).

P-bodies (PB) are cytoplasmic granules containing factors involved in mRNA decay and turnover that typically enlarge in stress conditions^18^. EDC3 is a scaffold protein for PB formation^26^, and its yeast homolog Edc3 forms condensates *in vitro*^27,28^. Unfortunately, the two EDC3 antibodies we tested only worked with methanol fixation, which is often incompatible with FP fluorescence. Therefore, we decided to use an antibody against the co-localizing PB marker protein DCP1 as a reference for FP-tagged EDC3 constructs (Figure S3A).

EDC3 fusion proteins were transiently expressed for 24 h and cells were stressed with sodium arsenite before fixation (Figure 3A). mCherry-tagging resulted in a marked reduction of PB numbers (Figure 3B), while other FP (EGFP, mEGFP, EYFP) and peptide tags (V5, myc) had minimal impact on PB number, area or IEC (Figure 3B, S3B). Similar to DDX3X, the influence of tagged EDC3 on the number and area of PB did not correlate with transient expression levels (Figure S3C/D).

**Figure 3:**
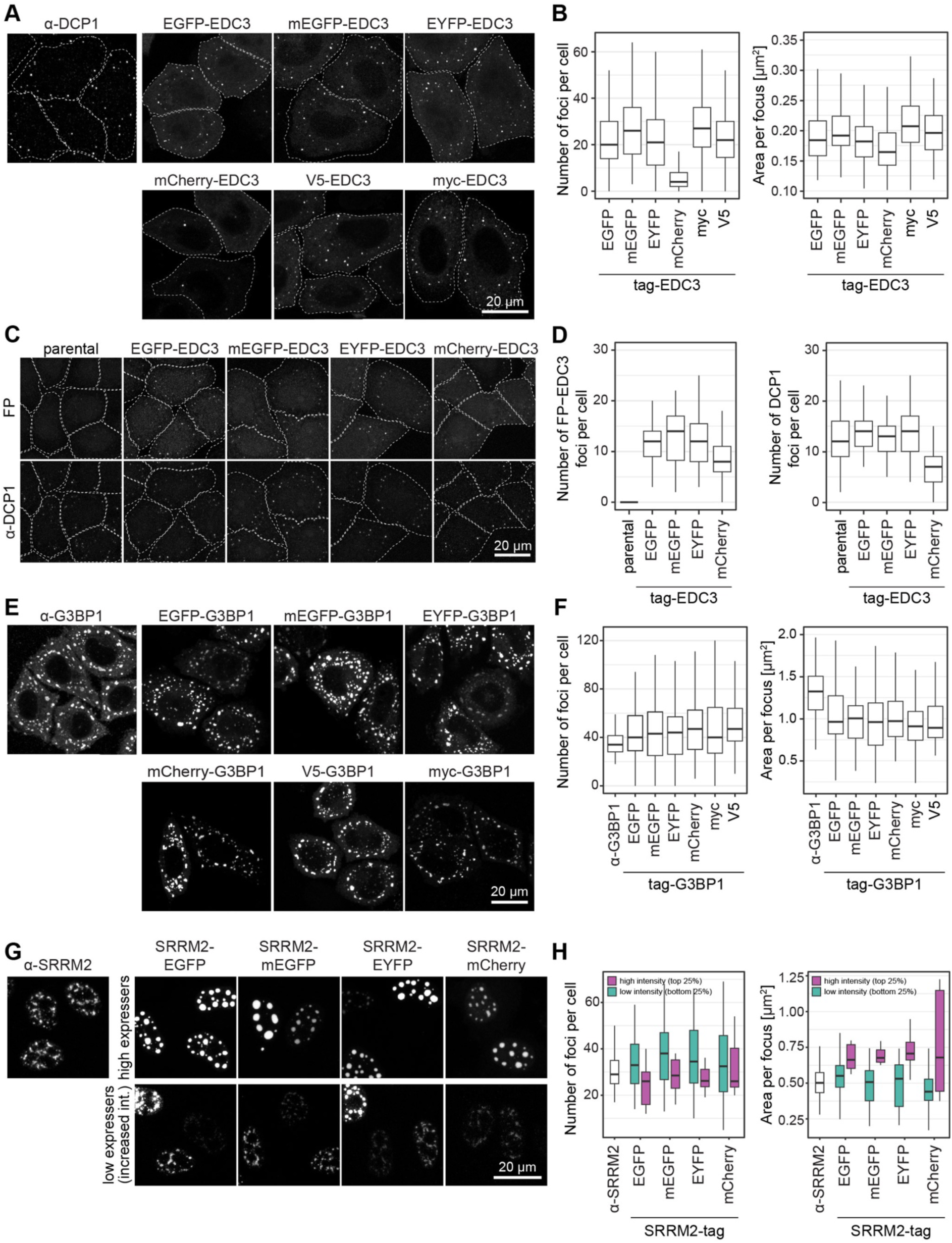
mCherry tagging reduces EDC3 foci but does not affect G3BP1 or SRRM2 foci formation Quantifications of foci include number of foci per cell and area per focus as indicated. (A) HeLa K cells were transiently transfected with EDC plasmids for 24 h, and stressed with 500 μM sodium arsenite for 30 min. V5-/myc-EDC3 were visualized by immunostaining with the respective antibodies. Untransfected cells were immunostained for DCP1 to visualize PB. (B) Quantification of (A); N = 3, n ≥ 110 cells. (C) Stable inducible HeLa cell lines were induced with doxycycline for 24 h to express the respective EDC3-tag constructs at endogenous level. Cells were stressed with 500 μM sodium arsenite for 30 min. For visualization of PB, cells were additionally immunostained for DCP1. (D) Quantification of (C). N = 2, n ≥ 55 cells. (E) HeLa K cells were transiently transfected with G3BP1 plasmids for 24 h and stressed with 500 μM sodium arsenite for 30 min. V5-/myc-G3BP1 were visualized by immunostaining with the respective antibodies. Untransfected cells were immunostained for G3BP1 as a control. (F) Quantification of (E); N = 3, n ≥ 90 cells. (G) HeLa K cells were transiently transfected with SRRM2 plasmids for 24 h and stressed with 500 μM sodium arsenite for 30 min. Untransfected cells were immunostained for SRRM2. (H) Quantification of (G) binned into low and high over expressing cells; N = 3, n ≥ 40 cells.

Stable cell lines expressing tagged EDC3 at endogenous levels confirmed that mCherry considerably reduced PB foci numbers (Figure 3C/D, S3E). Quantification of co-stained DCP1 foci revealed that mCherry-EDC3 expression drastically reduced DCP1 foci numbers, while other FP led to a slight increase (Figure 3D). This suggests that EDC3 tagging affects not only its own targeting to PB but also the recruitment of other proteins such as DCP1.

### Tagging G3BP1 or SRRM2 has minimal impact on stress granules and nuclear speckles, respectively

Tagging marker proteins does not always have a strong impact on CONDENSATE appearance, as illustrated by the following two examples. Condensation of G3BP1 was demonstrated to be crucial for SG formation ^29–31^. In our experiments, C-terminal fusion of G3BP1 to various FP and peptide tags did not notably affect the number, or IEC of G3BP1 SG foci (Figure 3E/F, S3F-H).

SRRM2 is a scaffold protein of nuclear speckles (NS), a nuclear CONDENSATE involved in splicing and storage of splicing factors^32,33^. Analysis of SRRM2 foci from transiently transfected tagged constructs revealed that SRRM2 expression levels had a greater impact on NS characteristics than the tags themselves (Figure 3G/H, S3I). Higher expression levels consistently resulted in larger, fewer NS, displaying distinct SON and SRRM2 sub-phases, consistent with the literature^34^.

In conclusion, while some CONDENSATE marker proteins are highly sensitive to tagging, others are minimally affected, even if they localize to the same CONDENSATE (e.g. DDX3X versus G3BP1). The impact of tagging varies across POI, and tagging can also influence other proteins recruited to the same organelle (e.g. EDC3 and DCP1). Furthermore, the results described above show that the effect of tagging can also depend on the expression levels.

### FP tagging of Pab1 alters SG formation in yeast, with mCherry strongly reducing granule formation

We wanted to investigate whether perturbations in fusion proteins also appear in yeast by tagging *S. cerevisiae* Pab1, the cytoplasmic poly(A) binding protein and a prominent SG marker, with various FP at the endogenous locus (Figure 4A). We observed striking differences in SG numbers depending on the FP used: ECFP, mTurquoise, and EGFP tags resulted in similar SG counts, EYFP doubled the count, mYPet caused a strong reduction, and mCherry or mScarlet-I nearly eliminated SG (Figure 4B, S4A/B). Similar but not identical trends were observed comparing two genetic backgrounds, W303 (Figure 4A/B) and BY (Figure S4A/B).

**Figure 4:**
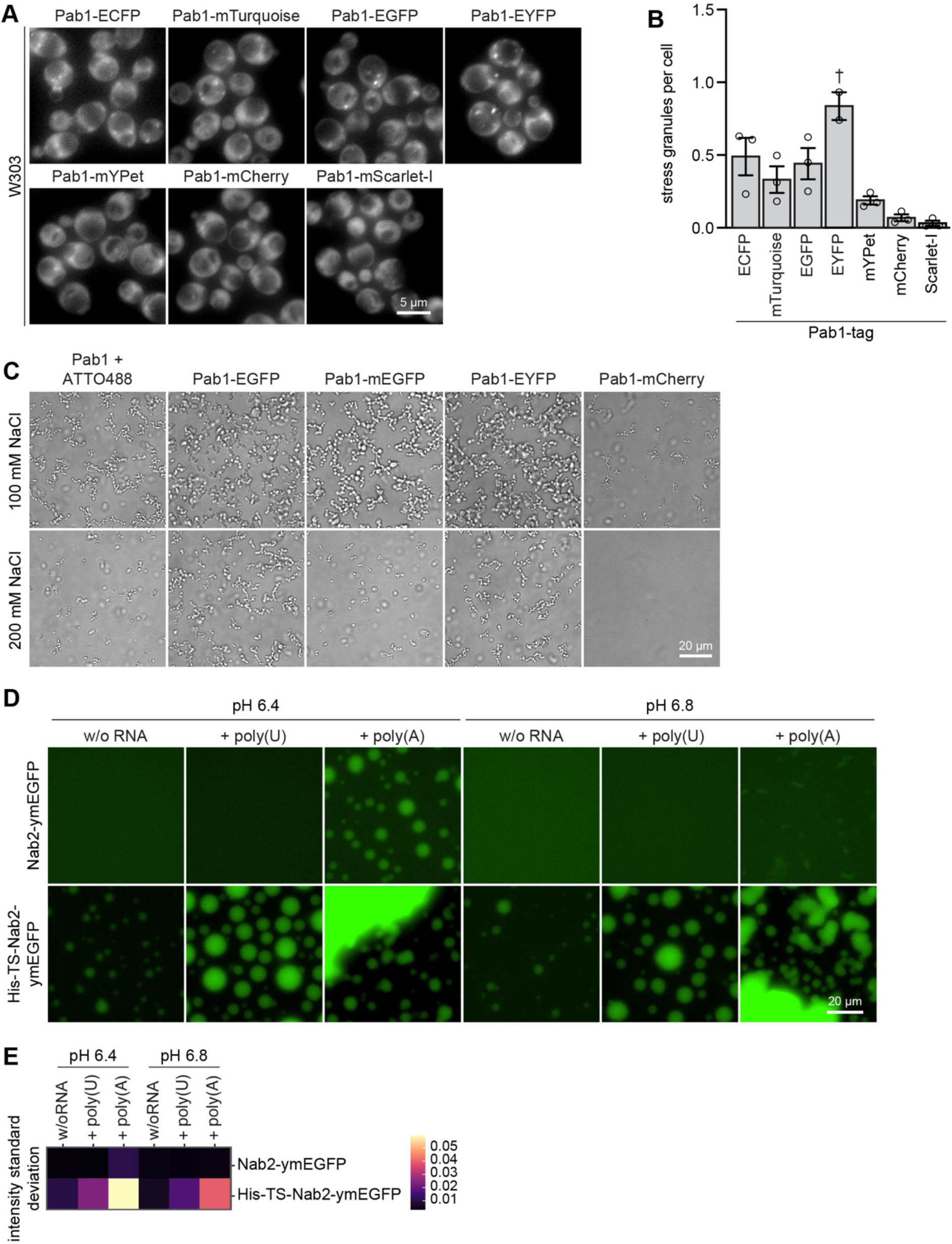
Tagging influences condensates of yeast polyA binding proteins *in vivo* and *in vitro* (A) *S. cerevisiae* W303 strains expressing tagged Pab1 were cultivated with 3% glycerol for 45 min prior to imaging to induce SG formation. (B) Quantification of SG per cell in (A), mean +/- SEM, N = 3, n ≥ 1500 cells per strain. For EYFP N = 2. (C) *In vitro* condensation assay with 15 µM Pab1 (FP-tagged or untagged + 1% ATTO488-Pab1 spike-in) in 25 mM NaOAc Buffer pH 5.0, at different salt concentrations (100 mM or 200 mM NaCl), 2 mM MgCl_2_. Incubation at 30°C for 15 min before imaging. N = 3. (D) Nab2 was purified and either used uncleaved as His10-TwinStrep(TS)-Nab2-ymEGFP or cleaved with Prescission protease to yield Nab2-ymEGFP. *In vitro* condensation assay with 20 µM protein in 25 mM sodium phosphate buffer pH 6.4 or 6.8, 50 mM NaCl, 2 mM MgCl_2_, 0.5 mg/mL BSA with 0.05 mg/mL poly(U) or poly(A) as indicated. Incubated at 20°C for 20 min before imaging. (E) Quantification of (D): image intensity standard deviation, N = 3.

Next, we examined the condensation behavior of Pab1 *in vitro* (Figure 4C, S4C). Consistent with previous findings, Pab1 formed hydrogel-like structures rather than roundish condensates (Figure 4C, S4D)^35,36^. Surprisingly, tagging substantially altered even these more aggregated structures: while EGFP, mEGFP, and EYFP-tagged Pab1 formed clusters similar in abundance to untagged+ATTO Pab1, though appearing somewhat larger and rounder, mCherry tagging drastically reduced Pab1 assemblies (Figure 4C, S4D).

### The His-TwinStrep affinity tag enhances Nab2 condensate formation and alters RNA binding specificity

*S. cerevisiae* Nab2, the nuclear poly(A)-binding protein, forms condensates *in vitro* and likely in cells^37^. FP tags were not tested in this case, but we found that a commonly used affinity tag, 6xHIS-TwinStrep (HisTS), can affect the RNA specificity of Nab2 condensate formation. HisTS-tagged Nab2-ymEGFP and cleaved Nab2-ymEGFP (Figure S4E/F) were tested for condensate formation in the presence of poly(A) RNA analog as their ‘natural’ substrate, and furthermore with poly(U) or without RNA (Figure 4D/E). While cleaved Nab2-ymEGFP formed condensates only with poly(A), HisTS-tagged Nab2 formed condensates under all tested conditions, including without RNA, and produced large assemblies with poly(A). This suggests that the HisTS tag adds nonspecific multivalency to Nab2, altering its RNA dependency and specificity. These findings highlight the importance of rigorously validating affinity tags in both *in vitro* and cellular assays.

### FP tags modulate condensation of *E. coli* CsdA *in vitro*

We extended our *in vitro* study to prokaryotic condensation-prone proteins by testing nine FP tags on the *Escherichia coli* DEAD-box ATPase CsdA. Initially, we examined the condensation behavior of CsdA with a larger and slightly different set of FP tags across different pH values (Figure 5A, S5C). Untagged+ATTO CsdA and CsdA fused with mEGFP, EBFP2, mKO2, and mNeonGreen formed condensates of similar size and PC (Figure 5A/B). In contrast, tagging with mYPet, mTurquoise2, and particularly mScarlet-I, mApple, and mCherry resulted in smaller, fewer condensates with lower PC, and at pH 8.0, these tags almost completely abolished condensation.

**Figure 5:**
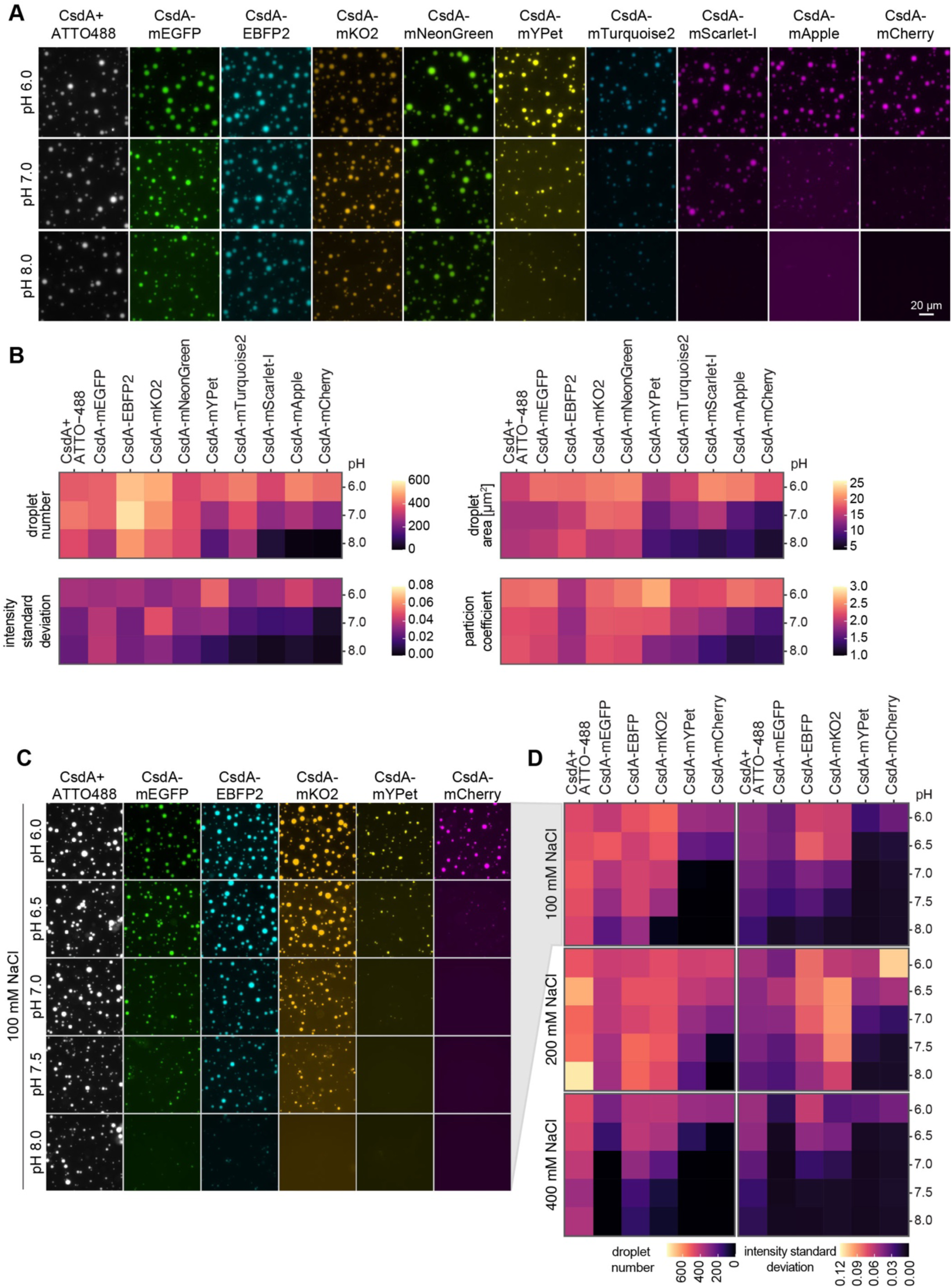
Fluorescent protein tags influence the condensation of *E. coli* CsdA *in vitro In vitro* condensation assays of recombinant CsdA (tagged or untagged + 1% ATTO488-CsdA spike-in) and quantifications (number of droplets per 0.1 mm^2^, droplet area, image intensity standard deviation and PC of N = 3). (A) 2 µM CsdA in 25 mM sodium phosphate buffer at the indicated pHs, 200 mM NaCl, 2 mM MgCl_2_, 0.5 mg/mL BSA, 0.05 mg/mL poly(U). Incubated at 25°C for 1 h before imaging. (B) Quantification of (A). (C) 2 µM CsdA in 25 mM sodium phosphate buffer at the indicated pHs, 100 mM NaCl, 2 mM MgCl_2_, 0.5 mg/mL BSA, 0.05 mg/mL poly(U). Incubated at 25°C for 1 h before imaging. (D) Quantification of (C and S5A).

A more detailed analysis of a subset of these FP tags examined their sensitivity to salt in addition to pH (Figure 5C/D, Figure S5A/B). While untagged+ATTO CsdA condensation was stable up to 400 mM salt across all pH values tested, FP-tagged CsdA showed increased sensitivity to salt, particularly at higher pH. Notably, some FP tags, such as mEGFP, mCherry, and mYPet, affected stability more than others like EBFP2 or mKO2. In summary, FP tags tend to reduce CsdA condensation, with the most pronounced effect at high salt and pH. CsdA-EBFP2 was closest to untagged+ATTO CsdA, while mCherry tagging significantly decreased condensate formation.

### *In vitro* condensation of FUS is sensitive to FP tagging

Next, we examined the *in vitro* condensation of FUS, an RNA binding protein implicated in amyotrophic lateral sclerosis (ALS) and frontotemporal dementia (FTD)^38,39^, and one of the most-studied proteins in condensation research. Tagging FUS with various FP revealed considerable differences in condensation dynamics (Figure 6A/B, S6A/B). At lower salt concentrations, FUS tagged with EGFP, mEGFP, or EYFP formed condensates similar to untagged+ATTO FUS. Under higher salt conditions (500/1000 mM NaCl), FUS-mEGFP remained comparable, whereas EYFP and EGFP tagged FUS showed increased droplet area and PC, especially at higher pH. This contrasts with untagged+ATTO FUS, where PC either decreased or remained constant with increasing pH. Once again, the mCherry tag strongly reduced condensation compared to other FUS variants. Overall, although FUS showed some sensitivity to FP tags, it was less affected than DDX3X, likely due to its more robust condensation under stringent conditions (e.g., 1000 mM NaCl). Based on our analysis, we recommend using the mEGFP tag for cellular studies of FUS as it most closely mimicked the behavior of untagged+ATTO FUS *in vitro*.

**Figure 6:**
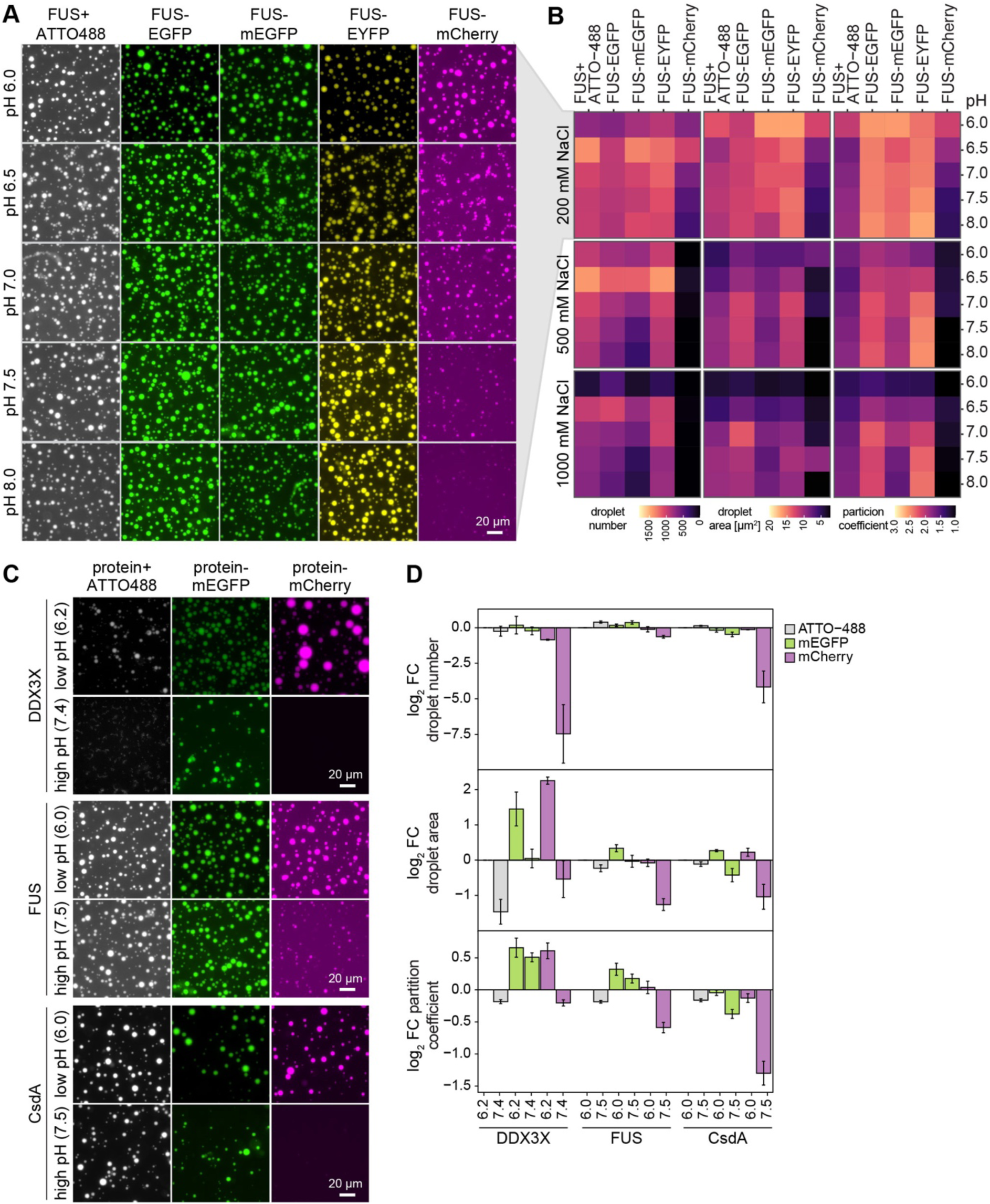
Fluorescent protein tags modulate FUS condensate behavior *in vitro* (A) *In vitro* condensation assay with 8 µM FUS (tagged or untagged + 2% ATTO488 FUS spike-in) in 25 mM sodium phosphate buffer at the indicated pHs, 200 mM NaCl, 2 mM MgCl_2_. Condensation was induced by addition of 10 µM His-3C-PreScission protease removing the MBP tag. Incubated at 25°C for 30 min before imaging. To account for pH-dependent FP intensity differences, we adjusted image brightness individually; corresponding images with consistent settings and DIC are in Figure S6A. (B) Quantification of (A and S6A): number of droplets per 0.1 mm^2^, droplet area and PC, N = 3. (C) Comparative assessment of DDX3X, FUS, and CsdA *in vitro* condensation when tagged with mEGFP or mCherry at 200 mM NaCl at pH 6.2/7.4 for DDX3X and pH 6.0/7.5 for FUS and CsdA as also displayed in Figures 2A, S5A and 6A. (D) Quantification of (C), log_2_ fold change (FC) of tagged protein against the respective untagged+ATTO pH 6.2/6.0 sample. N = 3.

### Comparison of *in vitro* condensation results for mEGFP and mCherry

To better visualize the effects of the two commonly used FP tags, mEGFP and mCherry, we compared their influence on the *in vitro* condensation of DDX3X, FUS, and CsdA at 200 mM salt and two pH levels (6.0/6.2 and 7.4/7.5) (Figure 6C/D). We used the quantifications from previous experiments (Figure 2, 5, 6) and normalized the changes relative to the untagged+ATTO samples at lower pH (Figure 6C/D). mEGFP tagging increased droplet area and PC for DDX3X, though still less than other FP tags (see Figure 1A), but it had minimal impact on FUS and CsdA condensation. In contrast, mCherry tagging had a more pronounced and complex effect. At low pH, mCherry induced large condensates with increased PC for DDX3X, while FUS and CsdA showed only moderate changes. However, at high pH, mCherry consistently impaired condensation for all three proteins. Notably, condensation of CsdA and FUS is more salt-resistant than DDX3X, suggesting that the DDX3X condensate interactions may overall be less stable and more easily influenced by FP tagging. Taken together, this highlights that mEGFP appears to be a relatively neutral tag for the POI tested here. In contrast, mCherry severely diminishes or even prevents the condensates of all tested POI, especially at higher pH.

### Phase coexistence simulations of fusion proteins

Given the substantial effects of FPs on the condensation of many proteins, we aimed to gain a deeper understanding of the underlying biophysical mechanisms of how protein tags may perturb phase equilibria. We build on previous work establishing a molecular grammar for phase separation^40^ and molecular simulation methods that can be used to relate protein sequence to propensities to undergo phase separation^41^. We performed direct-coexistence simulations on phase-separating proteins or intrinsically disordered regions (IDRs) fused to various FPs, using the CALVADOS 3 coarse-grained model, which we recently demonstrated can be used to predict the propensity of both IDRs and multidomain proteins to phase separate^41^. In a first experiment, we attached a diverse set of FP, namely ECFP, mEGFP, mNeonGreen, mCherry, mScarlet, Ruby3, to either the N- or C-terminus of full-length DDX3X and FUS, which consist of both folded domains and IDRs. For both proteins, we observed that FP-tagging influenced protein condensation, albeit to varying degrees, as visualized through equilibrium density profiles (Figure 7A, Figure S7A). DDX3X, with its complex amino acid composition and folded domain surface composition, was more strongly affected than FUS, which aligns with our biochemical data (Supplementary Table S2, Figure S8, Figure S9). While FP fusions at both the N- and C-termini of FUS generally exhibited similar condensation behavior, DDX3X showed noticeable differences in condensation depending on the tagging location (N- or C-terminus).

**Figure 7:**
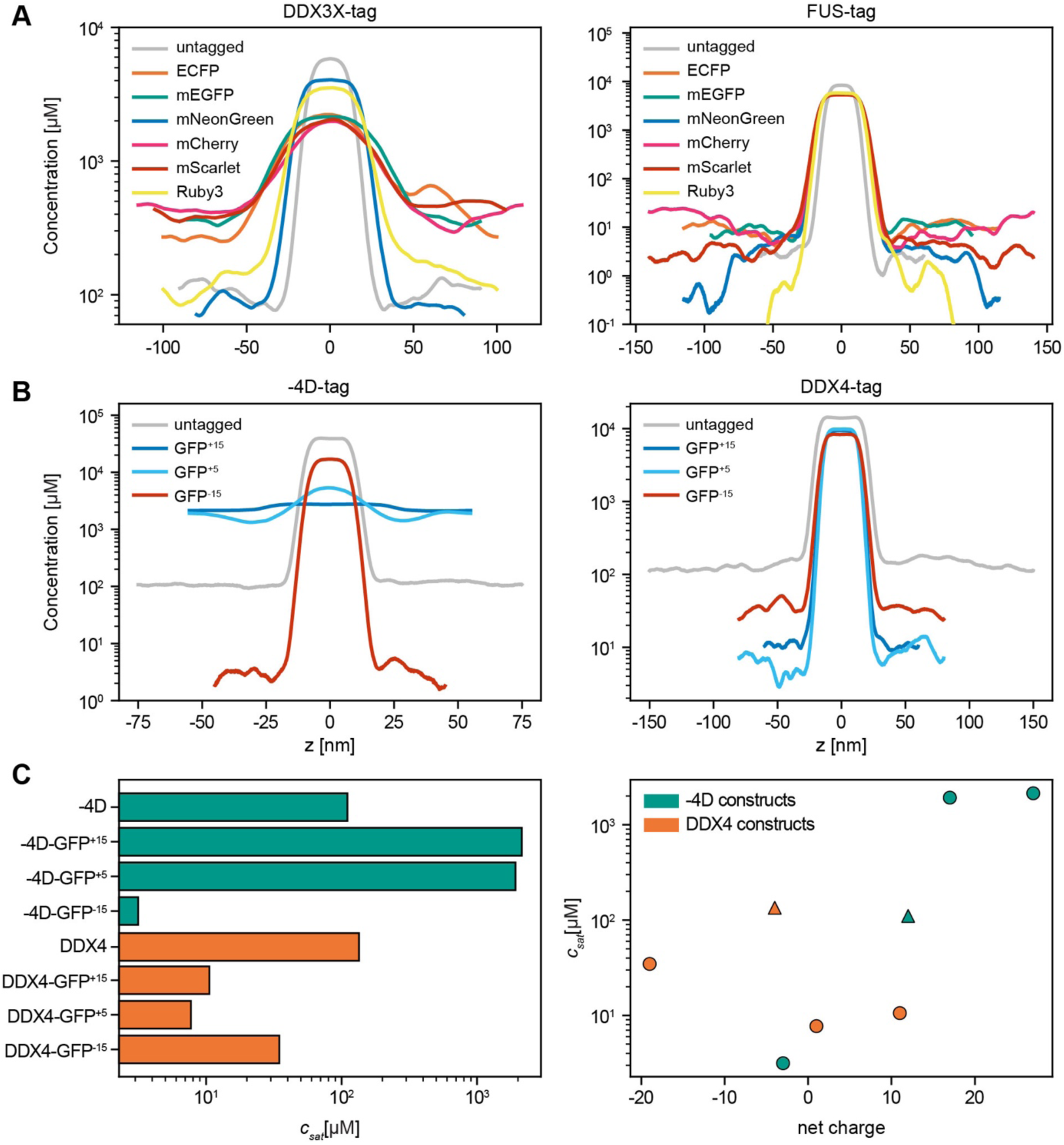
Net charge of fluorescent protein tags influences condensation behavior *in silico* Phase coexistence simulations were performed at 293 K with an ionic strength of 0.15 M using CALVADOS 3 (A) Equilibrium density profiles of six FPs C terminally fused with full-length DDX3X (left) and full-length FUS (right). (B) Equilibrium density profiles of the fully intrinsically disordered proteins-4D (left) and truncated DDX4 (right) C-terminally fused with three GFP variants with varying net charge. (C) Simulated saturation concentrations (c_sat_, left) and correlation between c_sat_ and net charge of for DDX4 (orange) or −4D (green) constructs with C-terminal tags. Circles represent tagged proteins, triangles untagged proteins.

Previous work has shown that protein net charge, together with other sequence properties, can affect the propensity to phase separate, so that proteins that have a net charge close to neutral appear to undergo homotypic phase separation more easily than charged proteins (in the absence of other effects)^21,42–44^. We therefore conduced a set of computational experiments to examine to what extent the net charge of the POI and the FP can help explain the effects of tagging. To do so, we shifted our focus to simpler systems consisting solely of IDR, specifically −4D^44^ where the four native Asp residues in the prion-like low-complexity domain of human hnRNPA1 have been substituted with Gly/Ser leaving the protein with a net charge of +12, and the isolated IDR of DDX4^45^ (with a net charge of −4), each fused to three GFP variants with differing net charges (GFP^-^^15^, GFP^+5^, and GFP^+^^15^). This allowed a clearer relationship to emerge between protein net charge and phase separation behavior. Notably, - 4D undergoes phase separation on its own despite its high net charge because of its high content of aromatic and arginine residues. However, the fusion of GFP^+^^15^ or GFP^+5^ (with overall charges of +15 and +5, respectively) abolished phase separation, likely due to electrostatic repulsion of the highly charged fusion protein (Figure 7B, Figure S7B, Figure S10, Figure S11). In contrast, attaching GFP^-^^15^ significantly promoted phase separation of −4D, presumably by a combined effect of diminishing the net charge of the fusion protein (nominal net charge of −3) and effects of GFP itself. Similarly, DDX4 IDR, with a net charge of −4, also undergoes phase separation on its own. The attachment of all three GFP charge variants enhanced condensation, presumably driven by additional interactions between the IDR and FP (Figure 7B/C, Figure S7B/C, Supplementary Table S3). In addition to the overall effect of adding GFP we find a systematic variation in the propensity to phase separate so that fusion of DDX4 IDR with GFP^+5^ (with a resulting net charge of +1) phase separates more strongly than the fusions to GFP^-^^15^ and GFP^+^^15^ (with net charges of −19 and +11, respectively). In summary, these results demonstrate that FP strongly influence phase separation, in part through their surface charges, though other effects also appear to play a role. These observations provide a starting point for developing or choosing FPs that minimally perturb condensate formation, though we note that cellular condensates have chemical environments that can be substantially more complex than the homotypic systems we have here examined.

## Discussion

Our study demonstrates that tagging a POI, especially with FP tags, can considerably influence its condensation properties *in vitro* and in cells. Given the many transient and weak interactions formed within a densely packed, multivalent condensate network, it is easy to perceive that tagging of their building blocks with proteins of considerable size and distinct surface properties may easily disrupt these interactions. Because the phase equilibria depend on interactions both within and outside a condensate, the tags may influence condensate formation due to interactions within the condensate or other cellular components. We find that these effects are not uniform, and that they vary depending on the tag, the POI and the physico-chemical environment. The properties of the tagged POI and the tag’s position relative to condensation-mediating residues likely contribute to these differences. This underscores the complex impact of protein tags on condensate and condensate formation. It is important to note that tagging can strongly influence the condensation of some POI while leaving others virtually unaffected. Even within the same condensate, different POIs can respond differently: for example, in SG, we found that DDX3X is affected by tagging while G3BP1 is not. It also appears that ‘scaffold proteins’ are not consistently affected: while DDX3X and EDC3 are impacted, G3BP1 and SRRM2 are not.

In general FP tags showed much stronger effects than peptide tags. EGFP and EYFP tend to promote condensate formation or even aggregation, likely due to their dimerization propensity or alternatively novel FP-mediated valency. Also, ligand-binding tags like Halo or SNAP, particularly when labeled with their respective dyes, can induce a more aggregated appearance. In contrast, mCherry and mScarlet-I strongly reduce condensate, and *in vitro* experiments reveal a strong pH-dependence for mCherry condensation. Most FP tags from the green wavelength spectrum used in this study (EBFP2, ECFP, EGFP, mEGFP, EYFP, mYPet, mTurquoise2) are derived from the same original protein, avGFP from *Aequorea victoria*, and overall behave very similarly. In contrast, the FP tags from the red wavelength spectrum (mCherry, mScarlet-I, mRuby3, mApple, mKO) are derived from distinct original FP and thus exhibit different tertiary structure (Figure S1A-C), explaining their more individualistic characteristics. These findings are consistent with recent studies showing that individual FP or affinity tags can disrupt condensation behavior in diverse systems, including viral factories, heterochromatin domains, and germline-specific granules^9–15,46,47^.

Given the complex effects observed for different FP tags, it is not surprising that changes do not appear to correlate directly with a single, easily determined biophysical property of the individual tags like size, isoelectric point or overall charge at a given pH (Supplemental Table S1). This suggests that multiple, more complex factors are involved. This is reflected by our coarse-grained simulations: attachment of FP often strongly modulate condensation, no matter whether the tag is attached at the N- or C-terminus of a POI. One particularly important factor seems to be the charge distribution on the protein surface^48^, which has also previously been shown to influence condensation^49^, as well as the addition of charged peptide tags^21,50^. Using coarse-grained simulations, we find that variations in the net charge of POI and GFP strongly influence to what extent and in which direction phase separation is modulated, consistent with electrostatic repulsion and attraction between the fusion proteins and the energetic cost of bringing in a charged protein to the dense phase. Since surface charge is, at least to some extent, pH-dependent, this may explain the strong pH sensitivity observed for certain FP like mCherry.

In this context, it is important to consider that at least in bacteria and yeast cells the intracellular pH of typically becomes more acidic in stress conditions, dropping from ∼7.4 to ∼6.4 ^51–53^, and that various condensates may establish complex gradients of pH and other ions^54,55^ that may in turn both be affected by protein tagging and modulate the effect of protein tagging. Consequently, the appearance of stress-induced CONDENSATE like PB or SG may be distorted if the POI is tagged with a FP that alters condensate structures in a pH-dependent manner.

### Implications for Future Research

The impact of protein tags on condensate appearance, as outlined in this study and described by others, carries significant implications for experimental design and interpretation. While there is no one-fits-all solution, we try to give some overall recommendations.

For experiments in cells, we recommend comparing tagged constructs to immunostaining of the endogenous protein when POI-specific antibodies are available. This provides a crucial reference for assessing how a tag may affect POI localization, recruitment to and ultimately its potential function within a biomolecular condensate. If no specific antibody exists, antibodies against other markers can help determine changes in overall condensate appearance. Additionally, benchmarking different tags against each other can offer further valuable insights. It is important to note that immunofluorescence staining requires fixation, which has been shown to alter the appearance of some condensates^4^. Thus, testing the effects of tags on condensation *in vitro* before transitioning to cellular studies can also provide valuable insights, although additional cellular parameters, including the condensate environment, may modulate the response.

Also, as is often the case in cellular work, overexpression of fusion proteins can artificially amplify or obscure condensation-related phenomena. Endogenous tagging, or stable cell lines where POI levels can be titrated to match endogenous protein levels — ideally paired with knock-down of the endogenous protein — can help reduce variability and minimize artifacts. This approach aligns with recent studies, which highlight that protein crowding and overexpression can mask or exaggerate the true phase-separation behavior of tagged proteins^13^.

For *in vitro* studies, we recommend avoiding FP tags as much as possible. Native recombinant untagged proteins can be observed using brightfield microscopy or, more quantitatively, with quantitative phase microscopy^23^ (REF as above). However, these approaches can be challenging or impractical in many experimental setups. In such cases, we suggest spiking in substoichiometric amounts (e.g. 1%) of POI that has been chemically labeled at defined sites (e.g. the N-terminus, or cysteines) with bright and pH stable dyes such as ATTO. In our experience, these dyes cause minimal interference across various experimental conditions.

Overall, FP choice is typically not straightforward. In the end, many factors, including the type of application, technical limitations of instruments, phototoxicity, and compatibility with other markers will guide the best choice. Careful tag selection, thorough validation with appropriate controls, and systematic analysis are key to mitigate tag-related issues and gain accurate insights into condensate behavior.

While these challenges pose a concern for many experiments, they could also become an opportunity: by switching FP tags, it may be possible to intentionally modulate condensate material properties^47^, shifting proteins to a more or less condensed, or even aggregated state, an approach that could offer valuable insights into the function of condensates.

## Supporting information

Suppmentary tables

## STAR Methods

### Key resource table

Key resource table:
| REAGENT or RESOURCE | SOURCE | IDENTIFIER |
| --- | --- | --- |
| **Antibodies** |  |  |
| anti-β-actin, C4 (mouse monoclonal) | Sigma | Cat# MAB1501;<br>RRID:AB_2223041 |
| anti-myc (mouse monoclonal) | Genetex | Cat# GTX29106;<br>RRID:AB_369669 |
| anti-V5 (mouse monoclonal) | Genetex | Cat# GTX635260; RRID:<br>AB_2888514 |
| anti-HA (rabbit polyclonal) | abcam | Cat# ab20084;<br>RRID:AB_445319 |
| anti-FLAG (mouse monoclonal) | Cell Signaling | Cat# 8146;<br>RRID:AB_10950495 |
| Ato488 FluoTag-X2 anti-ALFA (1G5) | Nanotag Biotech | Cat# N1502-At488-S; RRID:<br>AB_3075982 |
| anti-DDX3X (mouse monoclonal) | Abcam | Cat# ab196032; RRID:<br>AB_2910197 |
| anti-G3BP1 (rabbit polyclonal) | Genetex | Cat# GTX112191; RRID:<br>AB_11164802 |
| anti-EDC3 (mouse monoclonal) | Santa Cruz | Cat# sc-365024; RRID:<br>AB_10841422 |
| anti-DCP1A (rabbit, monoclonal) | Abcam | Cat# ab183709, RRID:<br>AB_3331658 |
| anti-SRRM2 ("SC35", mouse monoclonal) | Genetex | Cat# GTX11826;<br>RRID:AB_372954 |
| anti-Halo (rabbit, polyclonal) | Promega | Cat# G9281, RRID:AB_713650 |
| anti-SNAP (rabbit, polyclonal) | New England Biolabs | Cat# P9310S,<br>RRID:AB_10631145 |
| anti-SON (rabbit, polyclonal) | Genetex | Cat# GTX129778,<br>RRID:AB_2886086 |
| Alexa Fluor 488 goat anti-rabbit IgG (H+L) | Thermo Fisher Scientific | Cat# A-11008;<br>RRID:AB_143165 |
| Alexa Fluor 568 goat anti-rabbit IgG (H+L) | Thermo Fisher Scientific | Cat# A-11011;<br>RRID:AB_143157 |
| Alexa Fluor 647 goat anti-rabbit IgG (H+L) | Thermo Fisher Scientific | Cat# A-21244;<br>RRID:AB_2535812 |
| Alexa Fluor 488 goat anti-mouse IgG (H+L) | Thermo Fisher Scientific | Cat# A-11001;<br>RRID:AB_2534069 |
| **Bacterial and virus strains** |  |  |
| One Shot™ TOP10 Chemically Competent *E. coli* | Thermo Fisher Scientific | Cat# C404003 |
| Lemo21(DE2) Competent *E. coli* | New England Biolabs | Cat# C2528H |
| **Chemicals, peptides, and recombinant proteins** |  |  |
| FuGene | Promega | Cat# E5911 |
| SNAP-Surface 549 | NEB | Cat# S9112S |
| Halo TMR ligand | Promega | Cat# G8251 |
| Vectashield | Vector Laboratories | Cat# H-1200-10 |
| ATTO488 NHS-ester dyes | ATTO-TEC GmbH | Cat# AD 488-31 |
| PMSF Protease inhibitors | Carl Roth | Cat# 6367.3xxx |
| DNaseI | Roche | Cat# 10104159001 |
| RNaseA | Macherey-Nagel | Cat# 74050 |
| AcuteBand Pre-Stained Protein Ladder | Lubio | Cat# LU5001-2500 |
| Precision (3C) protease | GeneScript | Cat# Z02799 |
| Poly(A) | Sigma | Cat# P9403-25 mg |
| Poly(U) | Sigma | Cat# P9528-25 mg |
| Strep Tactin conjugated to HRP | iba | Cat# 2-1502-001 |
| **Experimental models: Cell lines** |  |  |
| HeLa K (HeLa Kyoto) | kind gift from Stefanie Jonas lab | RRID:CVCL_1922 |
| HeLa FRT | Gromeier lab; Kaiser et al., 2008 | N/A |
| HeLa FRT DDX3X-EGFP | this study | N/A |
| HeLa FRT DDX3X-mEGFP | this study | N/A |
| HeLa FRT DDX3X-EYFP | this study | N/A |
| HeLa FRT DDX3X-mCherry | this study | N/A |
| HeLa FRT DDX3X-myc | this study | N/A |
| HeLa FRT DDX3X-V5 | this study | N/A |
| HeLa FRT EGFP-EDC3 | this study | N/A |
| HeLa FRT mEGFP-EDC3 | this study | N/A |
| HeLa FRT EYFP-EDC3 | this study | N/A |
| HeLa FRT mCherry-EDC3 | this study | N/A |
| **Experimental models: Organisms/strains** |  |  |
| All *S.cerevisiae* strains generated in this study are listed in Supplementary table S5 | this study | N/A |
| **Recombinant DNA** |  |  |
| pETMCN_His10-TS-3C_DDX3X-mCherry | this study | N/A |
| pETMCN_His10-TS-3C_DDX3X-mEGFP(A206K) | this study | N/A |
| pETMCN_His10-TS-3C_DDX3X-EGFP | this study | N/A |
| pETMCN_His10-TS-3C_DDX3X-EYFP | this study | N/A |
| pETMCN_His10-TS-3C_DDX3X-ECFP | this study | N/A |
| pETMCN_His10-MBP-3C-DDX3X | this study | N/A |
| pETMCN_His10-TS-3C-MPB-3C_CsdA-YPET | this study | N/A |
| pETMCN_His10-TS-3C-MPB-3C_CsdA-mNeonGreen | this study | N/A |
| pETMCN_His10-TS-3C-MPB-3C_CsdA-mKO2 | this study | N/A |
| pETMCN_His10-TS-3C-MPB-3C_CsdA-mApple | this study | N/A |
| pETMCN_His10-TS-3C-MPB-3C_CsdA-mTurquoise | this study | N/A |
| pETMCN_His10-MPB-3C_CsdA-mCherry | this study | N/A |
| pETMCN_His10-TS-3C-MPB-3C_CsdA-mScarletl | this study | N/A |
| pETMCN_His10-TS-3C-MPB-3C_CsdA-EBFP2 | this study | N/A |
| pETMCN_His10-TS-MPB-3C_CsdA-mEGFP(A206K) | this study | N/A |
| pETMCN_His10-TS-3C_CsdA_untagged | this study | N/A |
| pETMCN_His10-TS-3C_mCherry | this study | N/A |
| pETMCN_His10-TS-3C_mEGFP(A206K) | this study | N/A |
| pETMCN_His10-TS-3C_EGFP | this study | N/A |
| pETMCN_His10-TS-3C_EYFP | this study | N/A |
| pETMCN_His10-TS-3C_ECFP | this study | N/A |
| pETMCN_His10-MBP-3C_Pab1-EYFP | this study | N/A |
| pETMCN_His10-MBP-3C_Pab1-mEGFP(A206K) | this study | N/A |
| pETMCN_His10-MBP-3C_Pab1-mCherry | this study | N/A |
| pETMCN_His10-MBP-3C_Pab1-EGFP | this study | N/A |
| pETMCN_His10-MBP-3C_Pab1_untagged | this study | N/A |
| pETMCN_His10MBP_3C_FUS-EYFP | this study | N/A |
| pETMCN_His10MBP_3C_FUS-ECFP | this study | N/A |
| pETMCN_His10MBP_3C_FUS-mEGFP(A206K) | this study | N/A |
| pETMCN_His10MBP_3C_FUS-mCherry | this study | N/A |
| pETMCN_His10MBP_3C_FUS-EGFP | this study | N/A |
| pETMCN_His10-MBP-3C_FUS_untagged | this study | N/A |
| pETMCN_His6-TS-3C_Nab2-ymEGFP | this study | N/A |
| pcDNA5_FRTTO_DDX3X-SNAP | this study | N/A |
| pcDNA5_FRTTO_DDX3X-YPet | this study | N/A |
| pcDNA5_FRTTO_DDX3X-V5 | this study | N/A |
| pcDNA5_FRTTO_DDX3X-mNeonGreen | this study | N/A |
| pcDNA5_FRTTO_DDX3X-ECFP | this study | N/A |
| pcDNA5_FRTTO_DDX3X-EYFP | this study | N/A |
| pcDNA5_FRTTO_DDX3X-Halo | this study | N/A |
| pcDNA5_FRTTO_DDX3X-mScarlet-I | this study | N/A |
| pcDNA5_FRTTO_DDX3X-FLAG | this study | N/A |
| pcDNA5_FRTTO_DDX3X-HA | this study | N/A |
| pcDNA5_FRTTO_DDX3X-ALFA | this study | N/A |
| pcDNA5_FRTTO_DDX3X-EGFP | this study | N/A |
| pcDNA5_FRTTO_DDX3X-mEGFP(A206K) | this study | N/A |
| pcDNA5_FRTTO_DDX3X-myc | this study | N/A |
| pcDNA5_FRTTO_DDX3X-mCherry | this study | N/A |
| pcDNA5_FRTTO_mCherry-G3BP1 | this study | N/A |
| pcDNA5_FRTTO_mEGFP(A206K)-G3BP1 | this study | N/A |
| pcDNA5_FRTTO_EYFP-G3BP1 | this study | N/A |
| pcDNA5_FRTTO_V5-G3BP1 | this study | N/A |
| pcDNA5_FRTTO_myc-G3BP1 | this study | N/A |
| pcDNA5_FRTTO_EGFP-G3BP1 | this study | N/A |
| pcDNA5_FRTTO_V5-EDC3 | this study | N/A |
| pcDNA5_FRTTO_EGFP-EDC3 | this study | N/A |
| pcDNA5_FRTTO_mEGFP(A206K)-EDC3 | this study | N/A |
| pcDNA5_FRTTO_mCherry-EDC3 | this study | N/A |
| pcDNA5_FRTTO_EYFP-EDC3 | this study | N/A |
| pcDNA5_FRTTO_myc-EDC3 | this study | N/A |
| pOG44 | Thermo Fisher Scientific | Cat# V600520 |
| pcDNA5_FRTTO_SRRM2-EYFP | this study | N/A |
| pcDNA5_FRTTO_SRRM2-mCherry | this study | N/A |
| pcDNA5_FRTTO_SRRM2-mEGFP(A206K) | this study | N/A |
| pcDNA5_FRTTO_SRRM2-EGFP | this study | N/A |
| **Software and algorithms** |  |  |
| OMERO | <sup>56</sup> | https://www.openmicroscopy.org/ |
| Cell Profiler | <sup>57</sup> | https://cellprofiler.org/ |
| Fiji | <sup>58</sup> | https://fiji.sc/; RRID: SCR_002285 |
| Fiji adaptiveThr function | ImageJ plugin by Quingzong Tseng | https://sites.google.com/site/qingzongtseng/imagejplugins |
| EasyFRAP | <sup>59</sup> | https://easyfrap.vimnet.upatras.gr/ |
| R studio | Posit team | https://posit.co/download/rstudio-desktop/ |
| Adobe Photoshop | Adobe | https://www.adobe.com/ |
| Adobe Illustrator | Adobe | https://www.adobe.com/ |
| AlfaFold | <sup>60</sup> |  |
| Prism 8 | GraphPad Software | [https://graphpad.com/features](https://graphpad.com/features;); RRID: SCR_002798 |
| FPbase | <sup>61</sup> | https://www.fpbases.org/ |
| Sedfit | <sup>62</sup> |  |
| **Other** |  |  |
| 384-well plate for condensation assays: black, optically clear flat-bottom, ultra-low-attachment-coated PhenoPlates | Revvity | Cat# 6057800 |
| 384 well plate for yeast imaging | Brooks | Cat# MGB101-1-2-LG-L |

### Resource availability

#### Lead Contact

Further information and requests for resources and reagents should be directed to and will be fulfilled by the lead contact, Maria Hondele.

#### Materials availability

All plasmids, yeast strains and human cell lines generated in this study are available upon request.

#### Data and code availability

No large-scale datasets have been generated in this study. The raw microscopy data that support the findings of this study are available from the lead contact upon reasonable request. This paper does not report original code. Additional information required to reanalyze the data reported in this paper is available from the lead contact upon request.

### Experimental model and study participant details

#### Human cell culture and cell lines

Human cell lines used in this study are listed in Supplementary Table S4. All cell lines were cultured at 37°C and 5% CO_2_ in DMEM supplemented with 10% fetal calf serum and 0.1 mg/mL penicillin/streptomycin. Tetracycline inducible cell lines were generated based on the FlpIn system (Invitrogen) allowing stable integration of pcDNA5-based constructs. Parental Hela cells containing an FRT site ^63^, were used to obtain monoclonal cell lines after selection with 0.3 mg/mL hygromycin B. All cell lines were tested negative for mycoplasma using PCR-based testing. Construct expression was induced with doxycycline.

#### Yeast cell construction and culturing

*Saccharomyces cerevisiae* strains used in this study are listed in Supplementary Table S5. Strains were constructed by transformation of a PCR amplification product with homology to the target site as described previously^64^. Cells were cultured in synthetic complete medium containing 2% D-glucose at 30°C.

#### Bacteria strains and culturing

*Escherichia coli* were cultured at 37°C in lysogeny broth (LB) medium with agitation at 150 - 200 rpm or on agar plates. Bacterial cultures carrying plasmids were supplemented with the corresponding antibiotic selection in the following concentrations; 50 μg/mL kanamycin, 100 µg/mL ampicillin or 30 µg/mL chloramphenicol. Cloning and plasmid amplification was done in *E. coli* Top10 and protein production carried out in *E. coli* Lemo21(DE2). Plasmids used in this study are listed in Supplementary Table S6.

### Method details

#### Plasmid construction

All proteins for expression in human cells were cloned into the pcDNA5 vector, with a GSGGSGG linker between the POI and the FP or peptide tag. Constructs for recombinant expression in *E. coli* and purification were cloned into pETMCN vectors, with the same GSGGSGG linker between POI and FP.

#### Transient transfection, drug treatment and fixation of human cells

Hela K cells grown on coverslips were transfected with plasmid DNA (listed in Supplementary Table S4) using FuGene transfection reagent. 24 h after transfection cells were fixed with 3% paraformaldehyde in PB for 15 min at room temperature, where indicated cells were treated with 500 µM sodium arsenite for 30 min prior to fixation to induce SG or PB formation. To label SNAP or Halo tagged proteins in cells, cells were incubated with either 2 µM SNAP-Surface 549 dye for 30 min before fixation or 200 nM TMR ligand for 15 min and washed three times with PB before fixation.

#### Immunostaining and microscopy of fixed human cells

After fixation with paraformaldehyde, cells were permeabilized with 0.1% Triton X-100, 0.02% SDS in PBS for 7 min. Alternatively cells were fixed with −20°C Methanol for 6 min at −20°C. Epitopes were then blocked with 2% BSA in PBS (BSA/PB) for 30 min. To stain the protein of interest, cells were next incubated with primary antibody (listed in key resources table) in BSA/PBS for 1 h. After three washes with PBS, cells were incubated with the respective secondary antibody diluted in BSA/PBS for 30 min. Following three washes with PBS, coverslips were mounted onto glass slides with Vectashield for confocal microscopy. Images were acquired using a Zeiss LSM700 Upright microscope with a 63x 1.4 NA oil DIC Plan-Apochromat objective. In all panels, representative maximum Z-projections are shown.

#### Immunoblot analysis

Cells were directly harvested with SDS sample buffer, boiled at 95°C for 5 min. Samples were separated on SDS-PAGE gels and proteins were transferred onto an activated PVDF membrane by wet blotting. The membrane was blocked with 4% milk in PBST for 30 min, before it was incubated with primary antibody diluted in 4% milk in PBT overnight at 4°C. After three washes for 5 min with PBT, the membrane was incubated in 4% milk containing the respective secondary antibody fused with HRP. Again, the membrane was washed three times and developed with ECL. The signal was detected using a Fusion (Vilber) imaging system.

#### Protein expression and purification

Recombinant proteins were expressed in *E. coli Lemo21(DE2)* by transforming chemically competent cells with the appropriate pETMCN-based expression plasmids under the respective antibiotic selection. Pre-cultures were grown overnight at 37°C in LB containing antibiotics and 1% D-glucose. Expression cultures were inoculated to yield an OD600 of 0.05 in Terrific Broth (TB) medium containing antibiotics and 1% D-glucose, grown at 37°C to an OD600 of 0.6-0.8 and induced with a final concentration of 200 mM IPTG. Cells were grown overnight at 18°C and harvested by centrifugation (5000 x g, 15 min, 4°C). Cell lysis was carried out in 30 mL lysis buffer (25 mM Tris-HCl pH 7.5, 1 M NaCl, 2 mM MgCl_2_, 10% glycerol (w/v), protease inhibitors, DNase and RNase) per 1 L of expression culture. After cell lysis by pressure homogenisation through an EmulsiFlex (Avestin) and / or sonication. Even if already lysed by Emulsiflex, sonication treatment can be critical to help with the removal of residual RNA after RNAse treatment (e.g. for FUS). Lysates were cleared by centrifugation (20’000 x g, 30 min, 4°C) and filtration through a 0.45 µm cut-off filter. His-tagged proteins were purified by immobilized metal affinity chromatography (IMAC) using self-made Ni^2+^ Sepharose columns, followed by 3C protease cleavage with simultaneous overnight dialysis. His10-TwinStrep-MBP tags and uncleaved proteins were removed by reverse IMAC. Proteins were further purified by size exclusion chromatography (SEC) using a 16/600 HiLoad Superdex 200 pg column (Cytiva) on an ÄKTA pure (GE Life Sciences) into the final storage buffer. IMAC, reverse IMAC and gel filtration fractions, as well as final protein pools were analyzed by SDS-PAGE and Coomassie staining, for size reference AcuteBand Pre-Stained Protein Ladder was loaded. Clean SEC fractions were pooled and concentrated using Amicon Ultra-4 centrifugal filters (Merck Millipore). Concentrated protein aliquots were snap-frozen in liquid nitrogen and stored at −80 °C. This general protein expression and purification protocol was applied to all DDX3X, CsdA, Pab1 and Nab2 constructs as well as the sole fluorescent protein constructs with small adaptations, i.e. by adjusting the buffer pH according to pI of the protein. In case of the FUS constructs, the 10xHis-MBP tag was not cleaved off, and subsequently reverse IMAC was not performed. Lysis methods, purification steps and storage buffer information for each individual recombinant protein are listed in Supplementary Table S7.

#### Chemical labeling of proteins

Untagged recombinant proteins were chemically labeled with ATTO488 NHS-ester dyes. To specifically label the primary amine of the N-terminus and not the lysines, labeling reactions were performed at pH 6.5 or lower, except for FUS, where labeling was performed at pH 7.5. If the protein storage buffer contained primary amines, buffer exchange into phosphate buffer was performed using Zeba Spin Desalting Columns, 7 K MWCO (Thermo Fisher Scientific) prior to reaction assembly. Protein and ATTO488 NHS-ester dye were mixed in a molar ratio of 1:4, followed by an incubation of 1 h at room temperature, protected from light. Using Zeba Spin Desalting Columns free dye was removed and labeled proteins were transferred back to the original protein storage buffer. Finally, labeled proteins were concentrated using Amicon Ultra-4 centrifugal filters (Merck Millipore) to 100-200 µM. Protein concentration and average degree of labeling was determined for each protein. Concentrated protein was aliquoted, snap-frozen in liquid nitrogen and stored at −80 °C. For microscopy-based assays labeled proteins were spiked-in, in a 1:100 molar ratio to the unlabeled protein, if not stated differently.

#### *In vitro* condensation assay

Condensation assays were carried out in 384-well, black, optically clear flat-bottom, ultra-low-attachment-coated PhenoPlates, following the procedure previously described by Hondele and colleagues^19^. In short, protein stocks were diluted using protein storage buffers to the 10-fold of the final assay concentration. For visualization of the untagged protein variants 1% of ATTO488 labeled protein was spiked-in the 10-fold concentrated protein stock. The 10-fold protein solution was transferred to the corner of a 384-well, where it was mixed in a ratio of 1:10 with corresponding assay master mixes, yielding final assay conditions as listed for the individual proteins in Supplementary Table S8. After reaction assembly plate was centrifuged (10 x g, 30 s, 25°C). The plate was incubated on the stage of the temperature-controlled microscope (25°C: DDX3X, FUS, CsdA and Nab2 or 30°C: Pab1). Reaction assembly and imaging were conducted at coordinated time intervals, with a constant total incubation time maintained across all tested fluorescent protein constructs to allow for direct comparison. Images were acquired on a temperature controlled, inverted Nikon Ti2 wide field microscope equipped with a 40x Plan Apo Lambda air objective NA 0.95, a Lumencor SPECTRA light source and a Hamamatsu ORCA-Fusion cMOS camera. All conditions were imaged using Differential Interference Contrast (DIC) microscopy and the appropriate fluorescence channels for each respective FP. Four images were taken in each of the three independent replicates. The microscope was controlled with the Nikon NIS Elements software and automated using the integrated JOBS system. Light intensity and exposure time were adjusted to the different fluorescent protein tags and maintained consistent within one experiment, and might differ between replicates. Images were processed by adjusting brightness and contrast using the open microscopy environment (OMERO). For each panel, representative images are shown.

#### Fluorescence recovery after photobleaching

Fluorescence recovery after photobleaching (FRAP) assays of *in vitro* condensates of DDX3X constructs (tagged or untagged + 1% ATTO488-DDX3X spike-in) were set up in 384-well, black, optically clear flat-bottom, ultra-low-attachment-coated PhenoPlates. Protein stock and condensation triggering master mix solutions were prepared identical to the *in vitro* condensation assay described above. Final assay conditions: 25 mM sodium phosphate buffer at the pH 6.2, 200 mM NaCl, 2 mM MgCl_2_, 0.5 mg/mL BSA, 0.05 mg/mL poly(U), 0.5 mM ATP, 0.5 mM DTT. The concentrated protein solutions were transferred to the corner of a 384-well, where it was mixed in a ratio of 1:10 with the corresponding assay master mix. Plate was centrifuged (10 x g, 60 s, 25°C), allowing reconstituted droplets to settle, followed by a total incubation of 1 h at room temperature prior FRAP acquisition. Photobleaching and image acquisition was performed on an inverted Olympus SpinD (CSU-W1) spinning disc microscope equipped with a RAPP UGA-42 Firefly 3L Photomanipulation/FRAP module, a 60x UPL APO oil objective NA 1.5 and a Hamamatsu ORCA-Fusion sCMOS camera. Circular region of interests (ROI) was bleached using a DL-473/100 laser. FRAP ROIs and laser intensity were controlled by SysCon2.0 software. The general FRAP experiment scheme included the acquisition of three pre-bleach frames, prior photo bleaching, followed by 180 post-bleach frames in a 1 s time interval. Standard deviation was assessed by comparing individual droplets within and across replicates. Images of DDX3X-mCherry were acquired using a 561 nm diode laser in combination with a 580-653 nm single bandpass filter. Images of DDX3X-mEGFP, DDX3X-EGFP, untagged DDX3X + 1% ATTO488-DDX3X spike-in and DDX3X-EYFP were acquired using a 488 nm diode laser in combination with a 500-550 nm single bandpass filter. Microscope and image acquisition was controlled with the Olympus cellSens software.

#### Analytical ultracentrifugation

Sedimentation velocity experiments were performed in 25 mM phosphate buffer pH 6.2 or 7.4, 150 mM NaCl, 1% glycerol, 1 mM MgCl_2_ and 0.5 mM β-mercaptoethanol, on 410 µL samples in double sector charcoal-epon centerpieces at 42,000 rpm and 20 °C using a Beckman XL-I analytical ultracentrifuge with the Beckman An-50 Ti rotor. Sedimentation was monitored during an overnight run using the interference detector with 900 scans and a nominal interscan delay of 1 min. The buffer densities (pH 6.2: 1.0105 g/ml, pH 7.4: 1.0098 g/ml) and viscosities (pH 6.2: 1.0214 centipoise, pH 7.4: 1.0443 centipoise) were measured at 20 °C using an Anton Paar DMA 4500M density meter and AMVn viscometer.

#### Microscopy of glycerol stressed yeast cells

Cells were inoculated from saturated overnight cultures into fresh synthetic media with the respective amino acid selection to OD_600_ 0.05-0.1, and grown at 30°C to OD_600_ 0.6–0.8. Cells were then transferred to a 384-well plate (Brooks, Matrical MGB101-1-2-LG-L) treated with Concanavalin A (stock solution of 1 mg/ml, air-dried), and spun down (100 x g, 1 min, 20°C). For acute glycerol stress, Pab1-tagged strains were washed three times in synthetic medium without glucose supplemented with final 3% glycerol directly in the imaging well, resuspended in the same medium and cultured for 45 min prior to imaging. All imaging experiments were performed at 30°C. Microscopy was performed using an inverted wide field microscope (Nikon Ti2) equipped with a Lumencor SPECTRA light source and a Hamamatsu ORCA-Fusion cMOS camera using a 60x Plan-Apo objective NA 1.4 and NIS Elements software.

#### Simulations

We generated fusion protein sequences by attaching tags to either the N-terminus or C-terminus of a POI. In this study we investigated four POI: two full-length proteins containing folded domains, DDX3X (FL_ DDX3X) and FUS (FL_FUS), and two intrinsically disordered proteins without folded domains, −4D^44^ and the isolated IDR of DDX4^45^. For FL_DDX3X and FL_FUS, we used AlphaFold to predict the structures of the fusion proteins. These predictions were then aligned with the published structures of the folded domains of DDX3X and FUS (catalytic core of DDX3X (PDB CODE: 5E7I), FUS-ZnF (PDB CODE: 6G99), FUS-RRM domain (PDB CODE: 2LA6)) to generate superimposed structures. The superimposed folded domains were then combined again with the IDRs from the AlphaFold predictions. Following relaxation of these structures, we used them as input for subsequent coarse-grained (CG) simulations. We used the following tags for simulations: ECFP, mEGFP, mNeonGreen, mCherry, mScarlet, Ruby3, GFP^+^^15^, GFP^+5^ and GFP^-15^ ^48^.

We applied the previously developed coarse-grained CALVADOS 3 model^41^ to perform direct-coexistence simulations in a cuboidal box. Each residue was represented as a single bead. C*_α_* positions were used for intrinsically disordered regions (IDRs) while center-of-mass position of residues were used for folded domains. The potentials of the model consist of a harmonic potential for chain connectivity, Ashbaugh-Hatch potential^65^ and Debye-Hückel potential^66^ for non-bonded interactions and another harmonic potential for restraining non-bonded residue pairs within the same folded domain. We chose to simulate 100 chains for each fusion protein. To avoid steric clashes of densely packed input structures, we selected the most compact conformation sampled by single-chain simulations with CALVADOS 3 and the corresponding CG representation as the initial conformation for each multi-domain protein (MDP) chain. Before production simulation, we performed equilibrium runs where an external force was used to push chains towards the box center so that a condensate could be formed. We then started production runs, saved frames every 0.125 ns and discarded the first 150 ns before analysis. The slab in every frame was centered in the box and the equilibrium density profile ρ(*z*) was calculated by taking the averaged densities over the trajectories as previously described^67^. Free energy of partitioning of fusion proteins were estimated as Δ*G*_*par*_ = *RT* ln *c*_*dil*_⁄*c*_*den*_ where *R* is the gas constant and *c*_*dil*_ and *c*_*den*_are the averaged concentrations of the dilute phase and dense phase (condensates), respectively.

### Quantification and statistical analysis

#### Quantification of foci in human cells

Maximum Z-projections were performed using Fiji software (version 2.14.0/1.54f). Cell profiler 2 (v 4.2.6) was employed to identify cells/nuclei and to segment DDX3X, EDC3, G3BP1, DCP1 and SRRM2 foci. After feature extraction, plots were generated using R studio.

#### Quantification of *in vitro* condensation assays

Cell profiler 2 (v 4.2.6) was used for droplet segmentation and feature extraction. Mean droplet number per 0.1 mm^2^, mean droplet area in µm^2^, mean partition coefficient (PC) and mean image intensity standard deviation was quantified and plotted using R Studio. Impact of fluorescent tagging with mEGFP and mCherry compared to untagged DDX3X, FUS and CsdA was assessed by calculating and plotting the mean log2 fold change (FP/untagged) of droplet number, droplet area, PC and image intensity standard deviation using R.

#### Quantification of analytical ultra centrifugation

The sedimentation velocity data were fitted to a diffusion-deconvoluted sedimentation coefficient distribution, c(s), using the software Sedfit 100 scans equally spaced in time and spanning the time period when the radial concentration profile was significantly changing were selected for analysis. A sedimentation coefficient range of 1-15 s was used with a resolution of 100 points. A single value of the frictional ratio (*f/f*_0_) for all sedimenting species, radialIy-invariant and time-invariant noise, baseline, meniscus and bottom position were all fitted.

#### Quantification of Fluorescence recovery after photobleaching

For analyzing the FRAP data individual droplets were stabilized using the StackReg plugin in Fiji software (version 2.14.0/1.54f) for image alignment. Next, images were analyzed using the easyFRAP software to quantify fluorescence recovery dynamics.

#### Quantification of foci in yeast cells

Images of yeast cells were processed using Fiji software (version 2.14.0/1.54f) and plots were generated in R studio. Granule quantification was performed using the Fiji adaptiveThr function (ImageJ plugin by Quingzong Tseng) with a weighted mean and strict thresholds to remove background and to isolate individual fluorescent foci.

#### Statistical analysis

Summary statistics including calculation of mean, median, N, standard deviation, standard error of the mean was performed using R. Sample size is indicated in the figure legend, where applicable.

## Acknowledgments

We would like to thank the other members of the Hondele lab, particularly Giulia Basile, Masroor Kahloon and Arturo Sanchez for helpful discussions and critical reading of the manuscript. We thank the imaging facility (IMCF) and the biophysics facility of the Biozentrum for their support on this project. M.G., and D.O. were funded by “Fellowships for Excellence” from the International PhD Program in Molecular Life Sciences of the Biozentrum, University of Basel. M.H. is funded by the Swiss National Science Foundation (PCEFP3_187052) and the European Research Council (ERC-ST2020 950262). K.L.-L. and F.C. are supported by the PRISM (Protein Interactions and Stability in Medicine and Genomics) centre funded by the Novo Nordisk Foundation (NNF18OC0033950, to K.L.-L.), and the China Scholarship Council (CSC 202206340019, to F.C.).

## Author contributions

K.D, D.O., K.L-L and M.H. conceived and designed the study. K.D., M.G., F.C., D.O., S.H., M.S., N.B. and T.S. performed the experiments and analyzed the data. K.D., M.G. and M.H. wrote the paper with input from the other authors.

## Declaration of interests

K.L.-L. holds stock options in and is a consultant for Peptone Ltd. The remaining authors declare no competing interests.

---

**Figure S1:**
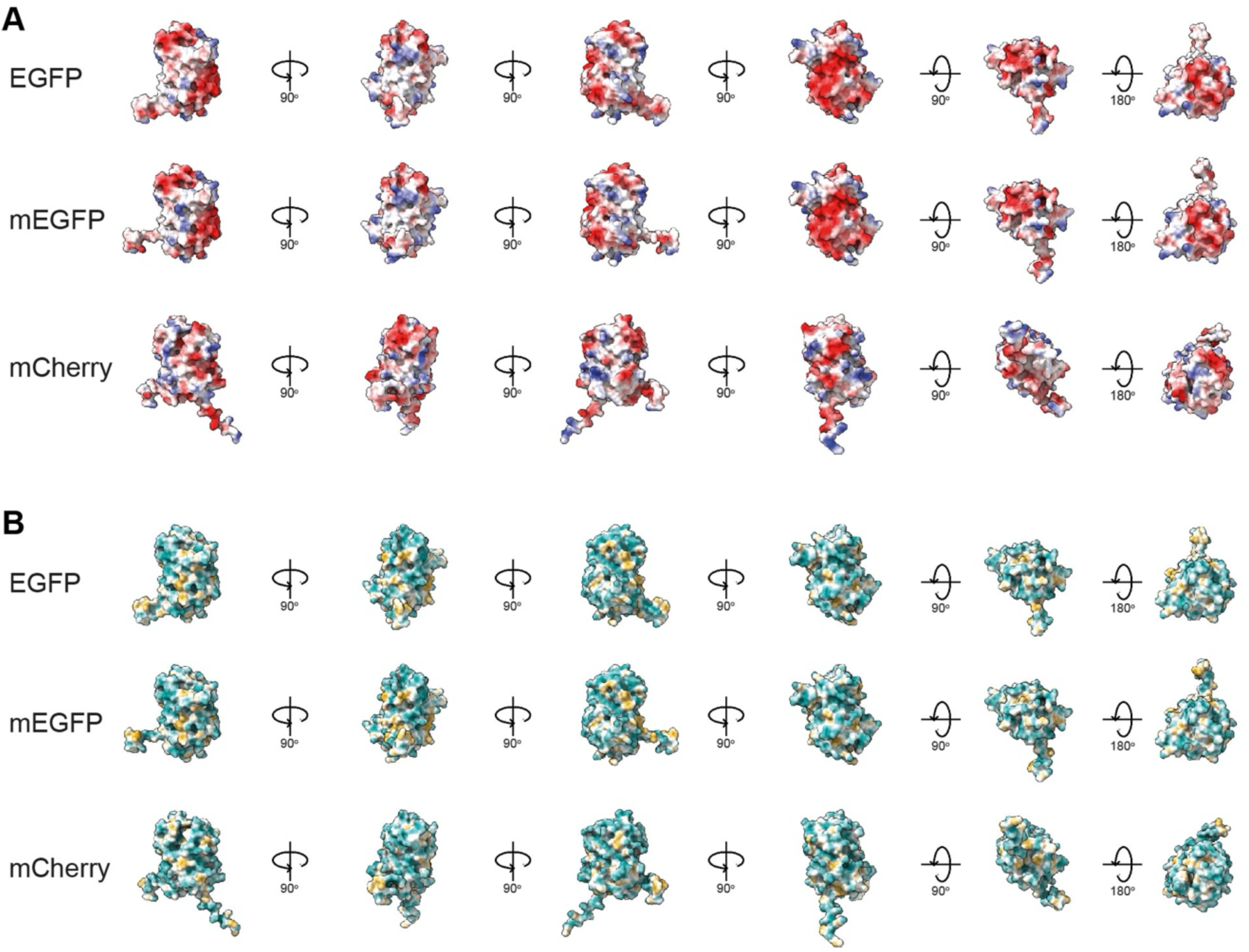
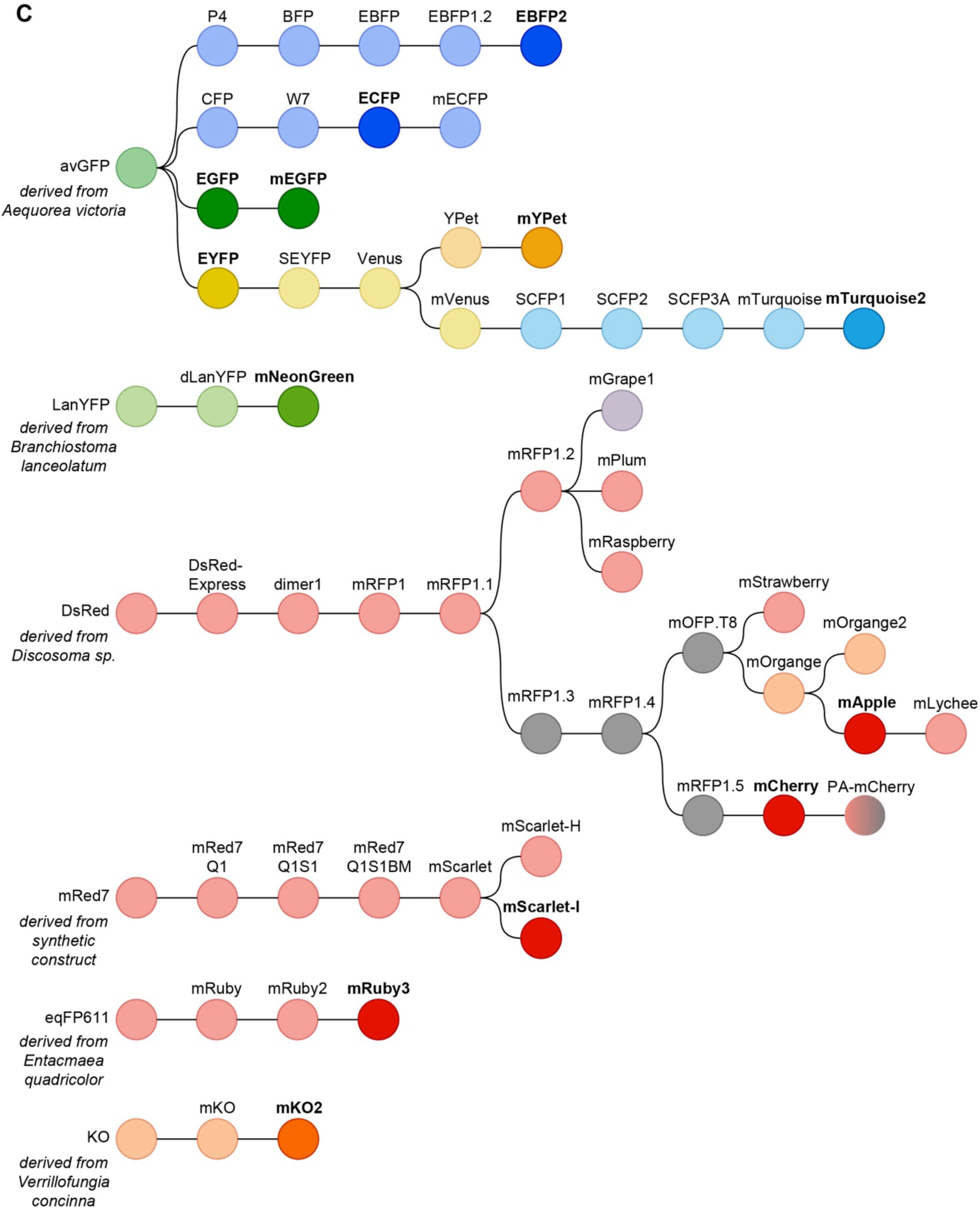
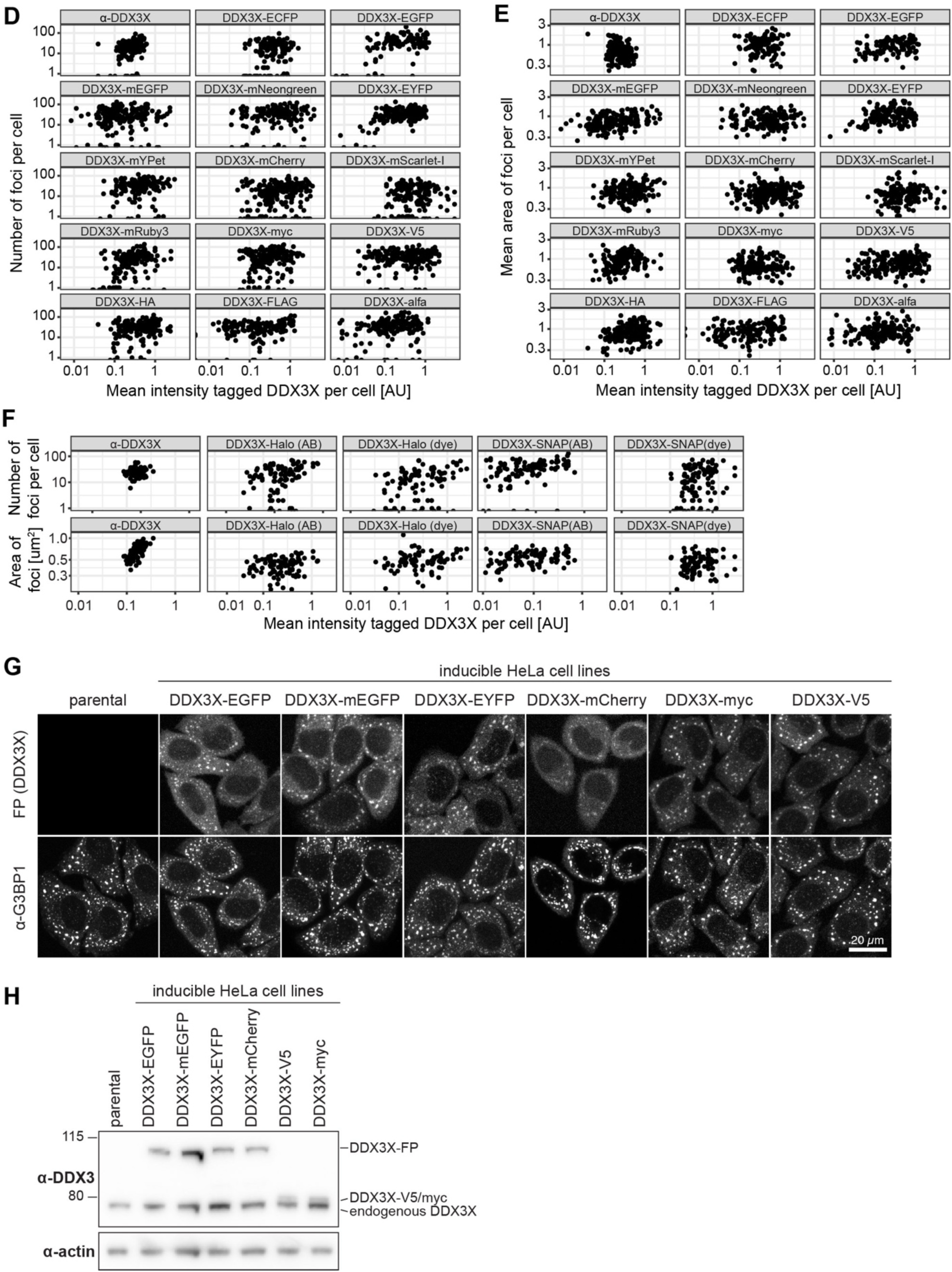
(A) Surface charge of selected FP from different angles, generated with AlphaFold v2.3. Positively charged patches are shown in red, negatively charged patches shown in blue. (B) Surface Hydrophobicity of selected FP from different angles, generated with AlphaFold v2.3. Hydrophobic patches are shown in beige, hydrophilic patches shown in cyan. (C) Overview of the evolutionary relationship among selected fluorescent proteins based on FPbase.org. Fluorescent protein tags used in this study are displayed in bold. (**D/E**) HeLa K cells were transiently transfected with DDX3X plasmids for 24 h, and stressed with 500 μM sodium arsenite for 30 min. Quantification of DDX3X foci as displayed in Figure 1A: number (D) and area (E) relative to DDX3X-tag expression level. N = 3, n ≥ 135 cells. (F) HeLa K cells were transiently transfected with DDX3X-Halo/SNAP plasmids for 24 h, and stressed with 500 μM sodium arsenite for 30 min. Halo and SNAP tags were either visualized by incubation with respective dyes (dye) or by immunostaining (AB). Quantification of DDX3X foci as displayed in Figure 1C: number and area relative to DDX3X-Halo/SNAP expression level. N = 3, n ≥ 90 cells. (G) Stable inducible HeLa cell lines were induced with doxycycline for 24 h to express the respective DDX3X-tag constructs at endogenous level. Cells were stressed with 500 μM sodium arsenite for 30 min before PFA fixation. For visualization of SG cells were additionally immunostained for G3BP1. Representative maximum Z-projections are shown. (H) DDX3X-tag expression levels of stable inducible HeLa cell lines in (**D**) were analyzed by immunoblotting with the indicated antibodies.

**Figure S2:**
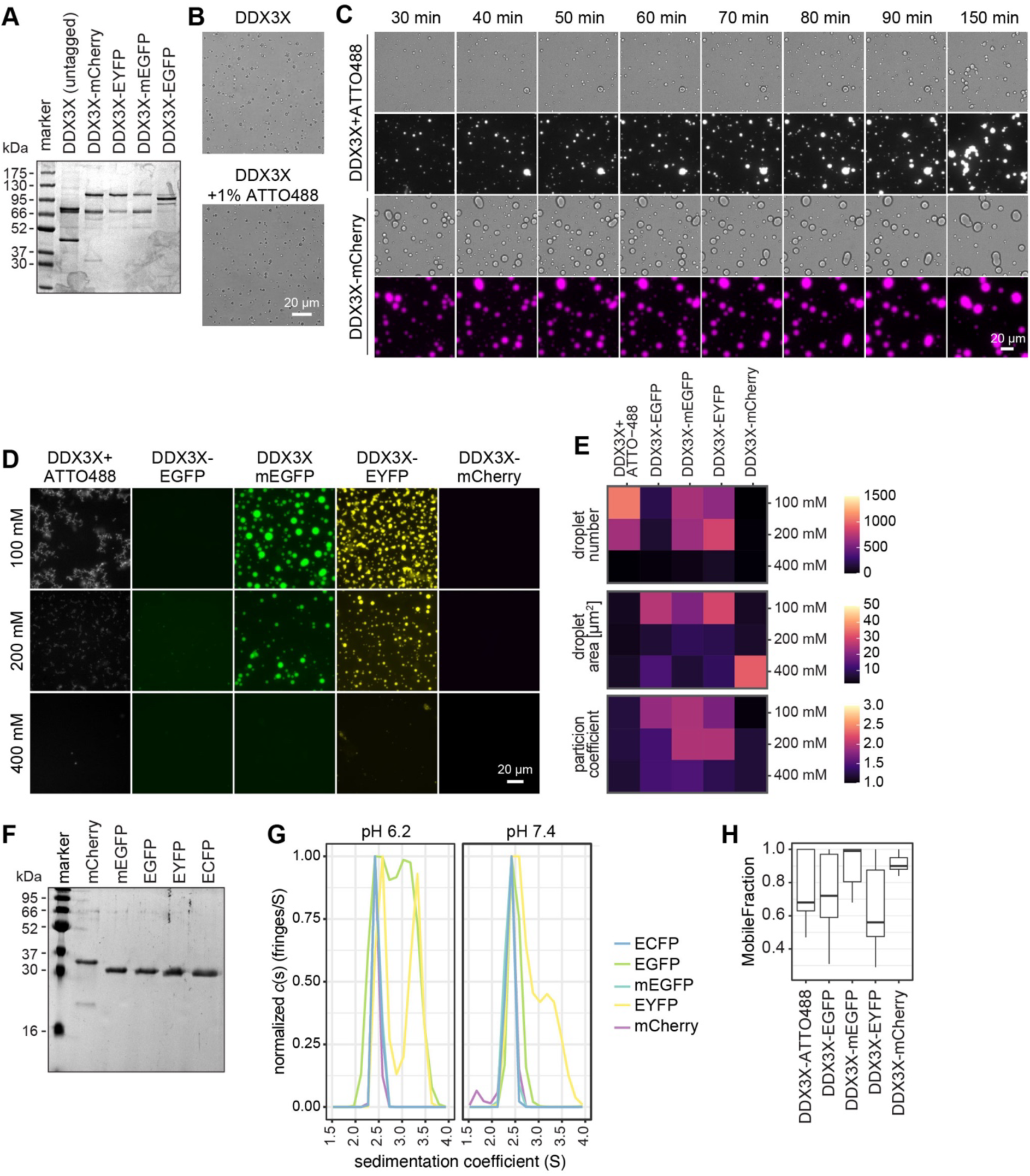
(A) Coomassie stained SDS PAGE with 0.5 µg of the respective DDX3X protein per lane. (B) *In vitro* condensation assay with 5 µM untagged DDX3X or 5 µM untagged DDX3X+1%ATTO-DDX3X spike-in in 25 mM sodium phosphate buffer at pH 6.2, 150 mM NaCl, 2 mM MgCl_2_, 0.5 mg/mL BSA, 0.05 mg/mL poly(U), 0.5 mM ATP, 0.5 mM DTT. Incubated at 25°C for 30 min before imaging. (C) Time series of *in vitro* condensation assay with 5 µM DDX3X + 1% ATTO488-DDX3X spike-in and DDX3X-mCherry in 25 mM sodium phosphate buffer at the pH 6.2, 150 mM NaCl, 2 mM MgCl_2_, 0.5 mg/mL BSA, 0.05 mg/mL poly(U), 0.5 mM ATP, 0.05 mM DTT. Incubated at 25°C for the indicated time before imaging. (D) *In vitro* condensation assay with 5 µM DDX3X (tagged or untagged + 1% ATTO488-DDX3X spike-in) at the pH 7.4, 100, 200 or 400 mM NaCl, 2 mM MgCl_2_, 0.5 mg/mL BSA, 0.05 mg/mL poly(U), 0.5 mM ATP, 0.5 mM DTT. Incubated at 25°C for 30 min before imaging. (E) Quantification of *in vitro* condensation assay in (D): number of condensates per 0.1 mm^2^, droplet area and PC, N = 3. (F) Coomassie stained SDS PAGE with 0.5 µg of the respective FP per lane. (G) Analytical ultracentrifugation of 0.5 mg/mL of the respective fluorescent protein in 25 mM phosphate buffer pH 6.2 or 7.4, 150 mM NaCl, 50 mM KCl, 1% glycerol, 1 mM MgCl_2_ and 0.5 mM β-mercaptoethanol. Values were normalized to the highest value for each construct. (H) FRAP of *in vitro* DDX3X condensates of similar size. 5 µM DDX3X in 25 mM sodium phosphate buffer at the pH 6.2, 100 mM NaCl, 2 mM MgCl_2_, 0.5 mg/mL BSA, 0.05 mg/mL poly(U), 0.5 mM ATP, 0.5 mM DTT. Condensates were matured for 1 h at 25°C before FRAP. Mobile fraction was analyzed. N = 3, n ≥ 23 condensates.

**Figure S3:**
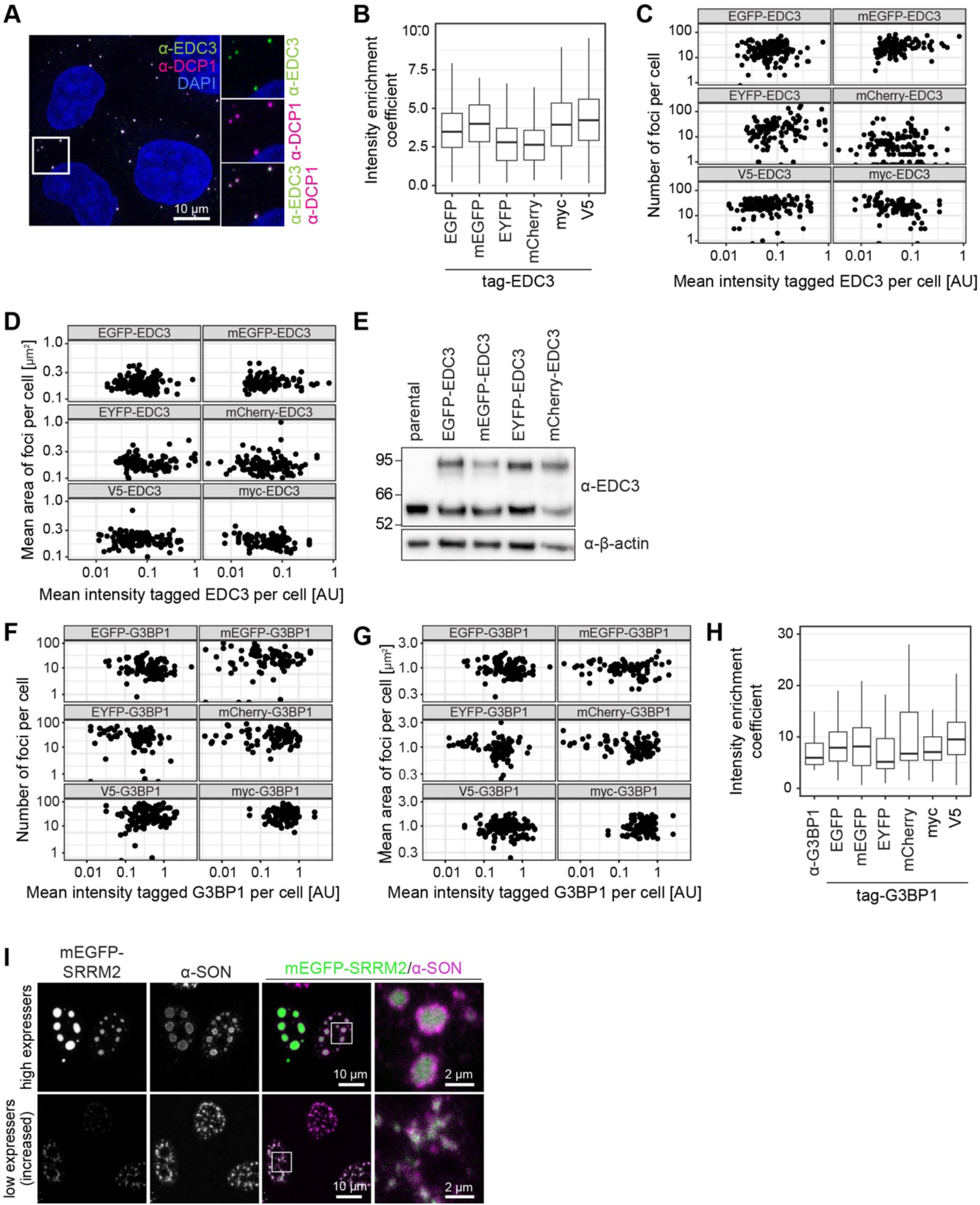
(A) Stressed HeLa K cells (500 μM sodium arsenite, 30 min) were fixed with methanol and immunostained for EDC3 and DCP1 with respective antibodies. (**B/C/D**) HeLa K cells were transiently transfected with EDC plasmids for 24 h, and stressed with 500 μM sodium arsenite for 30 min. V5-/myc-EDC3 were visualized by immunostaining with the respective antibodies. Quantification of EDC3 foci of cells in Figure 3A: intensity enrichment coefficient (median intensity per focus/ median intensity of the cell) (B), number (C) and area (D) relative to the cellular expression level. N = 3, n ≥ 110. (**E**) Tag-EDC3 expression levels of stable inducible HeLa cell lines from Figure 3C were analyzed by immunoblotting with the indicated antibodies. (**F/G/H**) HeLa K cells were transiently transfected with G3BP1 plasmids for 24 h and stressed with 500 μM sodium arsenite for 30 min. V5-/myc-G3BP1 were visualized by immunostaining with the respective antibodies. Untransfected cells were immunostained for G3BP1 as a control. Quantification of G3BP1 foci as displayed in Figure 3E: Number (F) and area (G) relative to the cellular expression level, intensity enrichment coefficient (median intensity per focus/ median intensity of the cell). (H). (**I**) HeLa K cells were transiently transfected with a plasmid for expression of mEGFP-SRRM2. After 24 h, cells were stressed with 500 μM sodium arsenite for 30 min. For visualization of nuclear speckles cells were additionally immunostained for SON. Representative centre plane images are shown.

**Figure S4:**
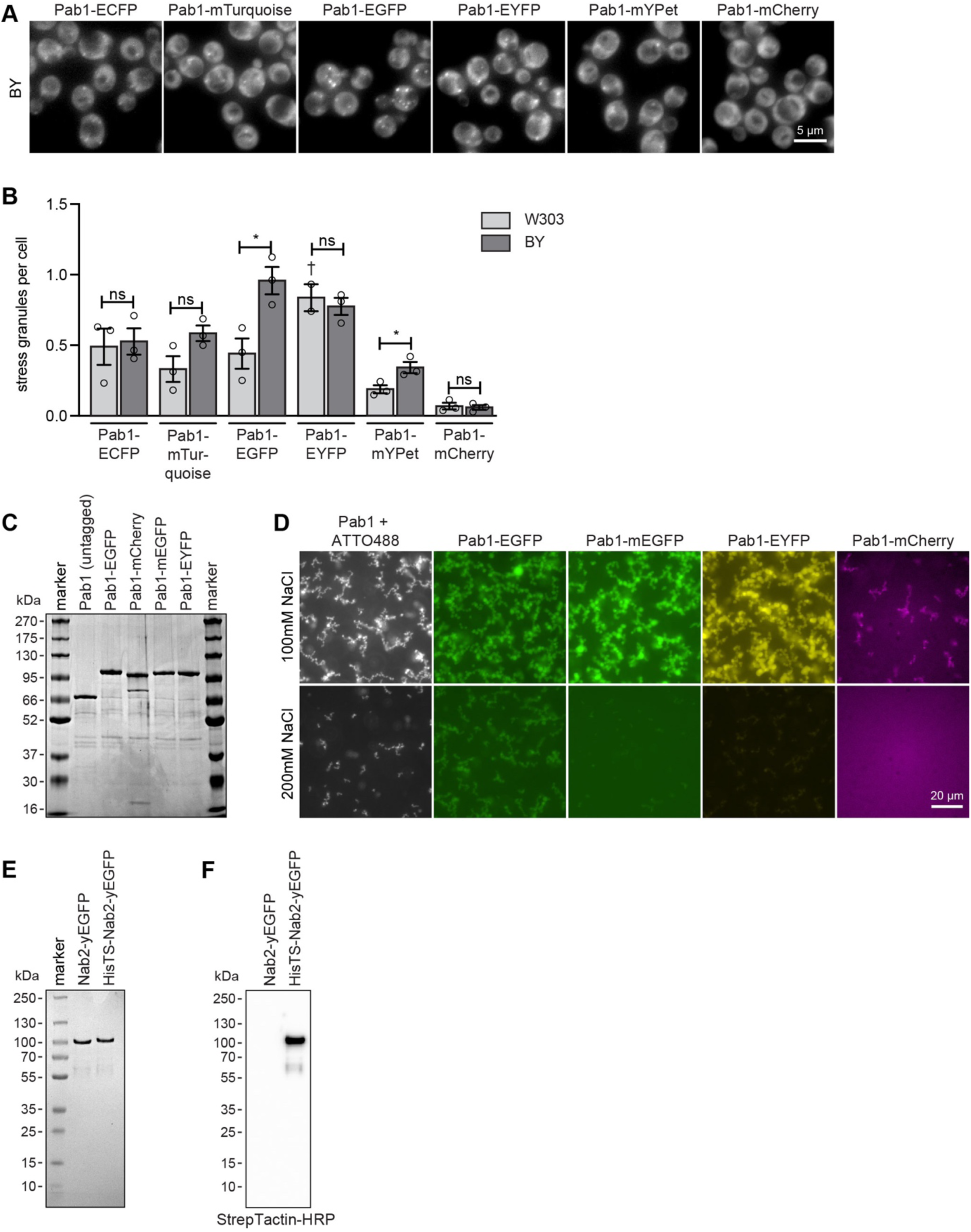
(A) *S. cerevisiae* BY strains expressing Pab1 fused with the indicated FP tag were cultivated with 3% glycerol for 45 min prior to imaging to induce SG formation. Representative maximum Z-projections are shown. (B) Quantification of SG per cell in (A) and Figure 4A, mean +/- SEM, N = 3, n ≥ 1500 cells per strain. ns: non-significant, *p≤0.05 in unpaired t-test W303 against BY. (C) Coomassie stained SDS PAGE with 0.5 µg of the respective Pab1 protein per lane. (D) FP channel images corresponding to DIC images in Figure 4C. (E) Coomassie stained SDS PAGE with 0.5 µg of the respective Nab2 protein per lane. (F) Western Blot with 0.5 µg of the respective Nab2 protein per lane, stained with Strep Tactin conjugated with horse radish peroxidase (HRP).

**Figure S5:**
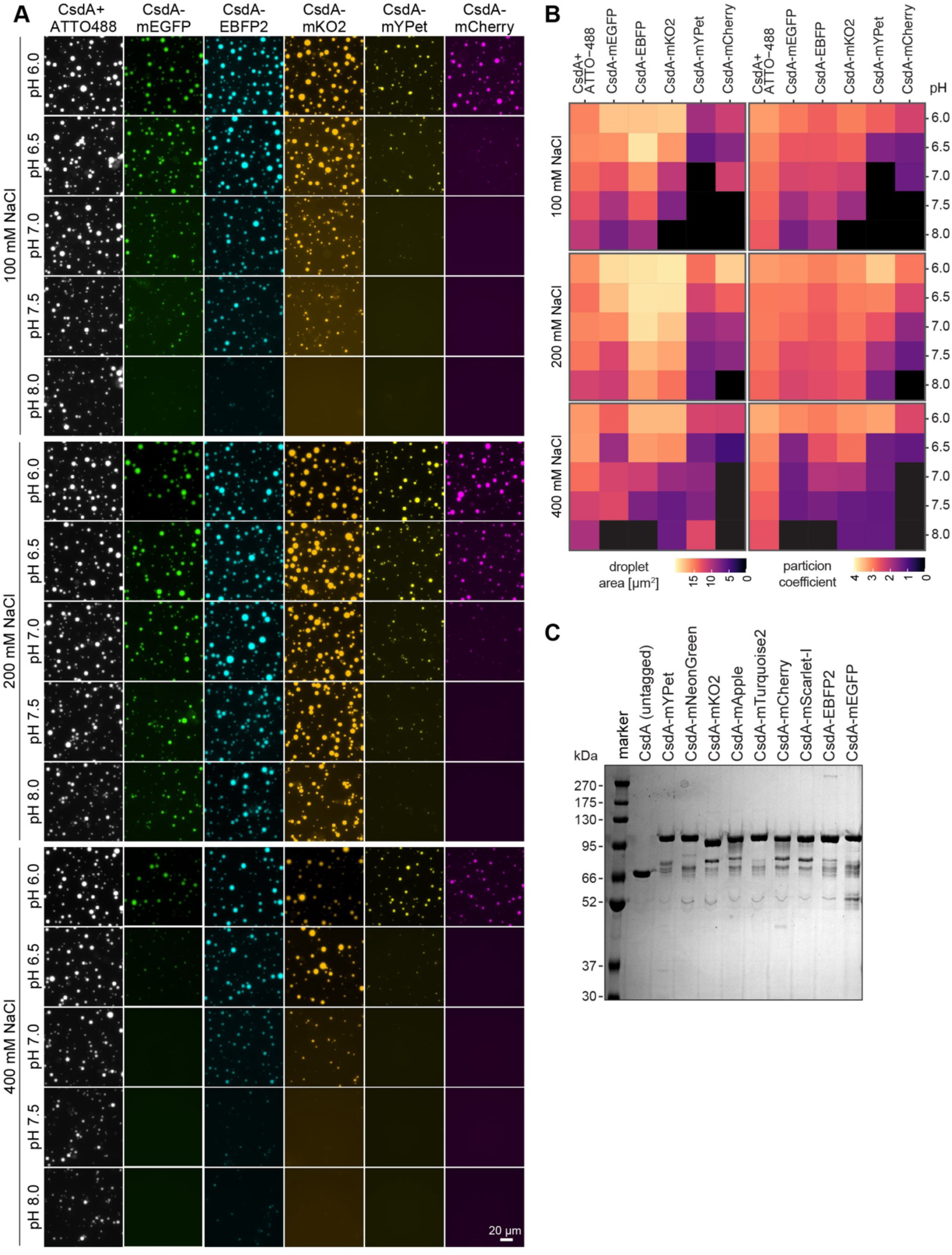
(A) *In vitro* condensation assay with 2 µM CsdA (tagged or untagged + 1% ATTO488-CsdA spike-in) in 25 mM sodium phosphate buffer at the indicated pHs and NaCl concentrations, 2 mM MgCl_2_, 0.5 mg/mL BSA, 0.05 mg/mL poly(U). Incubated at 25°C for 1 h before imaging. Representative images are shown. (B) Further quantification of *in vitro* condensation assay in (A), in addition to Figure 5C, N = 3. (C) Coomassie stained SDS PAGE with 0.5 µg of indicated CsdA protein per lane.

**Figure S6:**
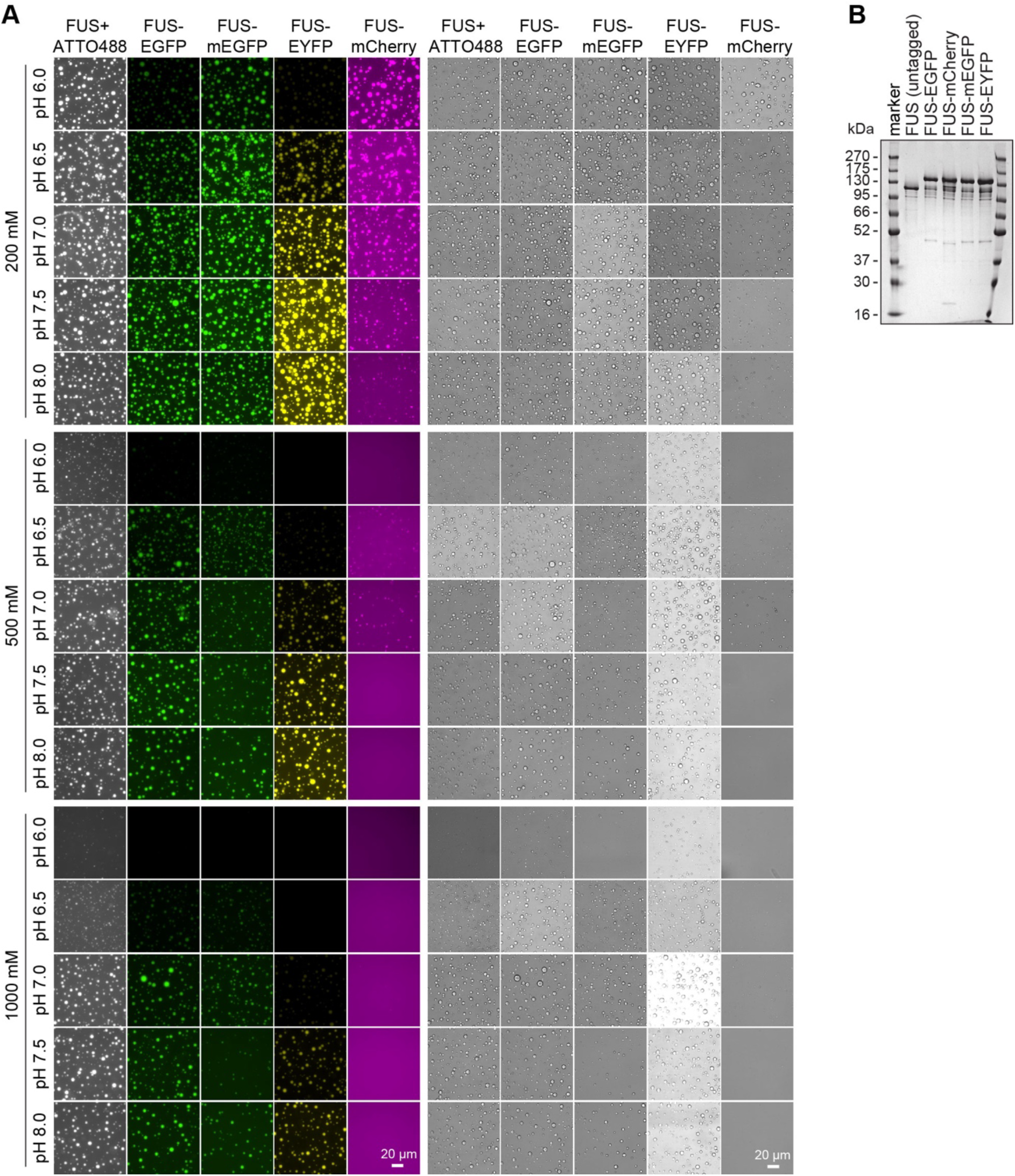
(A) *In vitro* condensation assay with 8 µM FUS (tagged or untagged + 2% ATTO488-FUS spike-in) in 25 mM sodium phosphate buffer at the indicated pHs and NaCl concentrations, 2 mM MgCl_2_. Condensation was induced by addition of 10 µM His-3C protease removing the solubility tag. Incubated at 25°C for 30 min before imaging. Representative images are shown. (B) Coomassie stained SDS PAGE with 0.5 µg of indicated FUS protein per lane.

**Figure S7:**
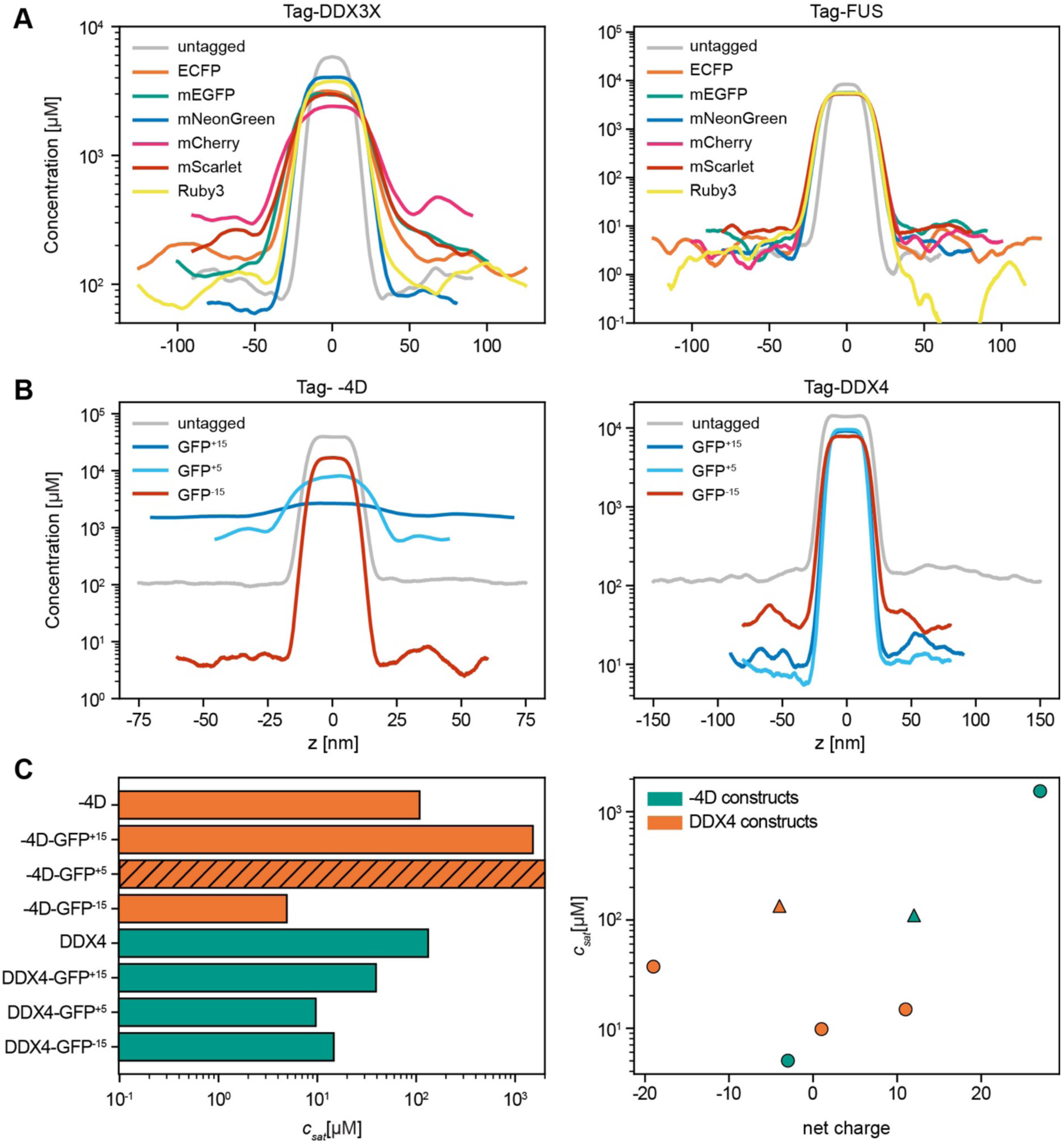
Phase coexistence simulations were performed at 293 K with an ionic strength of 0.15 M using CALVADOS 3 (A) Equilibrium density profiles of six FPs N-terminally fused with full-length DDX3X (left) and full-length FUS (right). (B) Equilibrium density profiles of the fully intrinsically disordered proteins −4D (left) and truncated DDX4 (right) N-terminally fused with three GFP variants with varying net charge. (C) Simulated saturation concentrations (c_sat_, left) and correlation between c_sat_ and net charge of for DDX4 (orange) or −4D (green) constructs with N-terminal tags. Circles represent tagged proteins, triangles untagged proteins. Hatched bar: We did not see a stable condensed phase.

**Figure S8:**
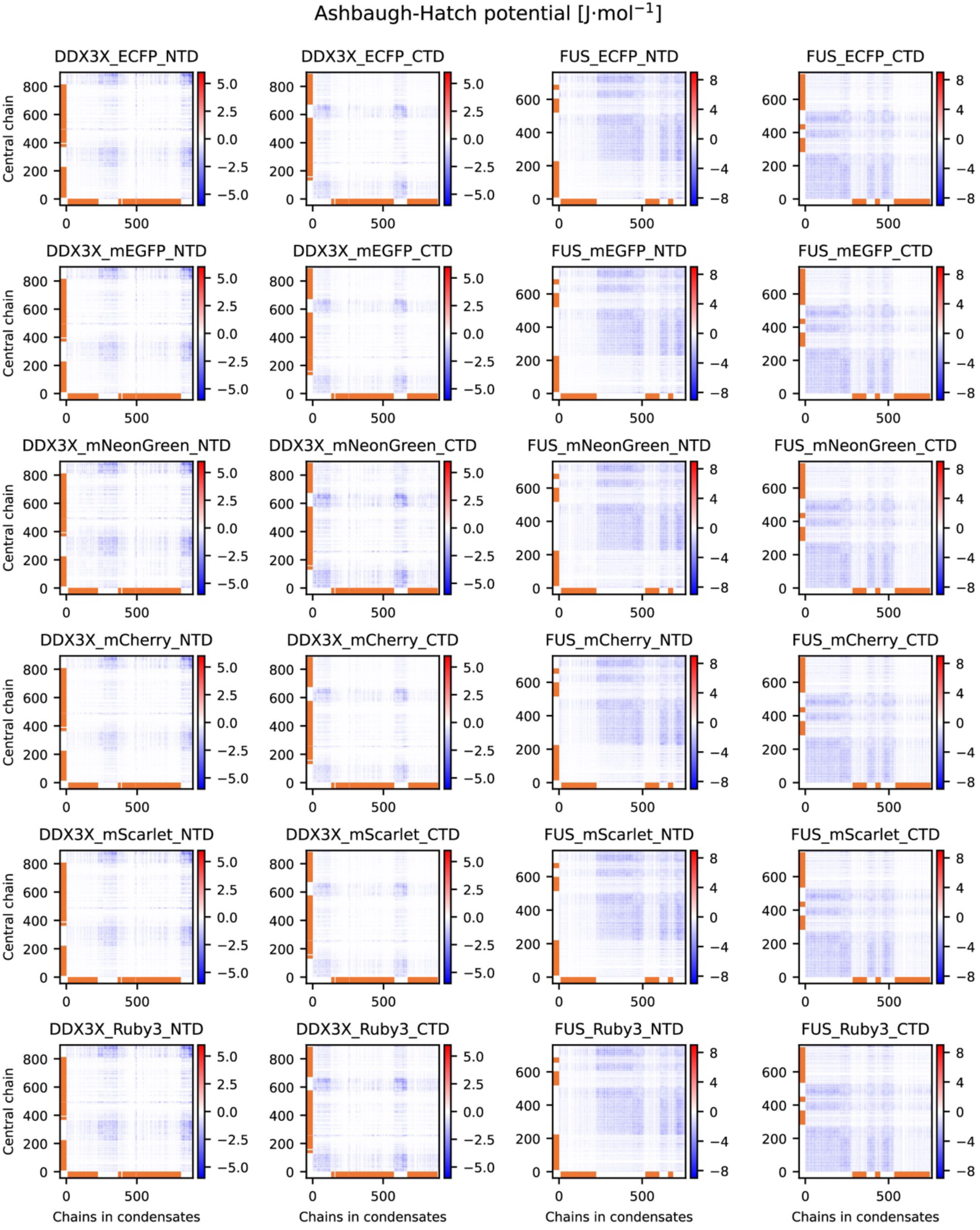
Averaged residue-pair interaction energies (the Ashbaugh-Hatch potential in forcefield) between the most central chain and the rest of the condensate for DDX3X and FUS with six different florescent proteins. The orange lines indicate folded parts.

**Figure S9:**
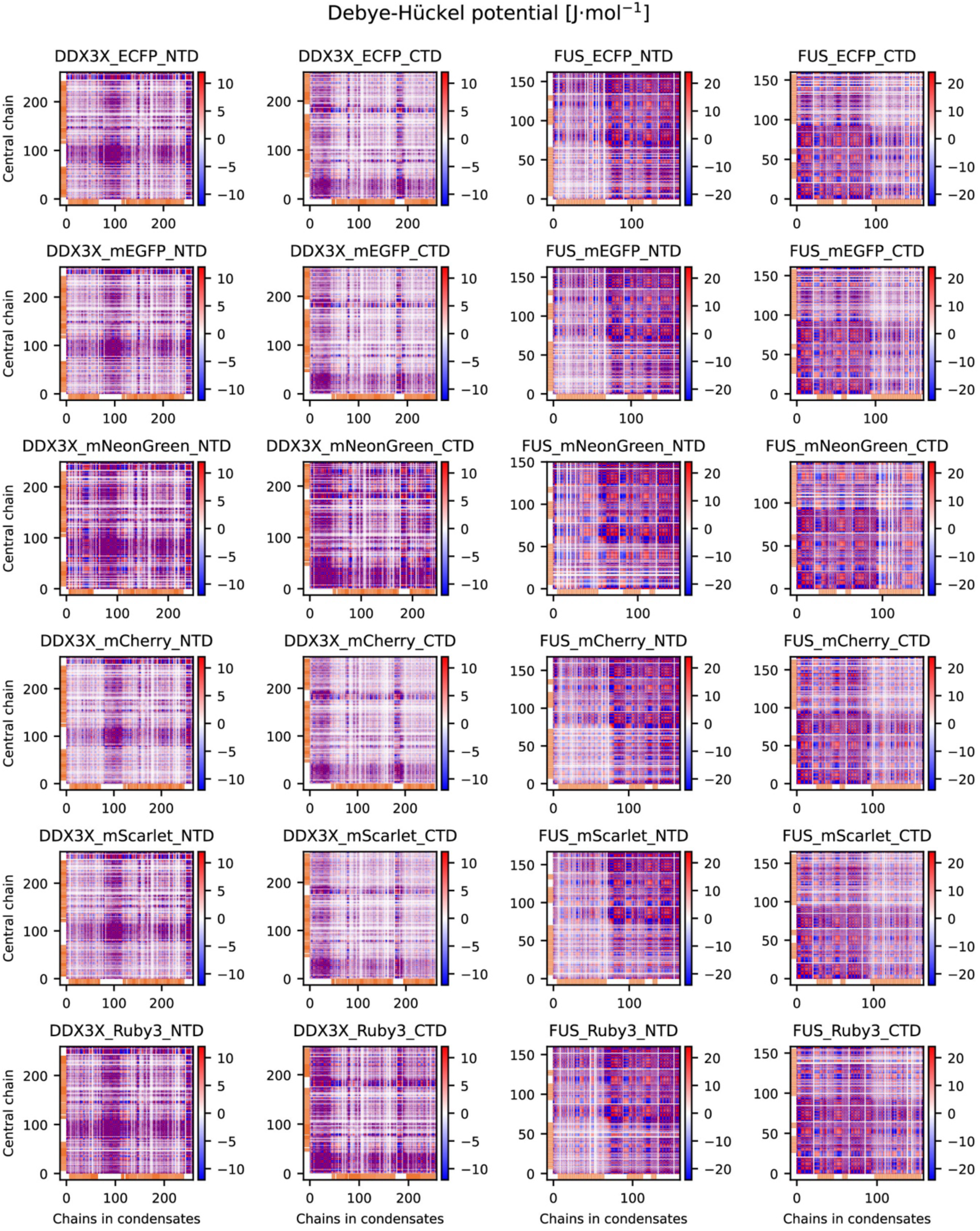
Averaged residue-pair electrostatic interaction energies (the Debye-Hückelpotential in forcefield) between the most central chain and the rest of the condensate for DDX3X and FUS with six different florescent proteins. Only the charged residues are shown. The orange lines indicate folded parts.

**Figure S10:**
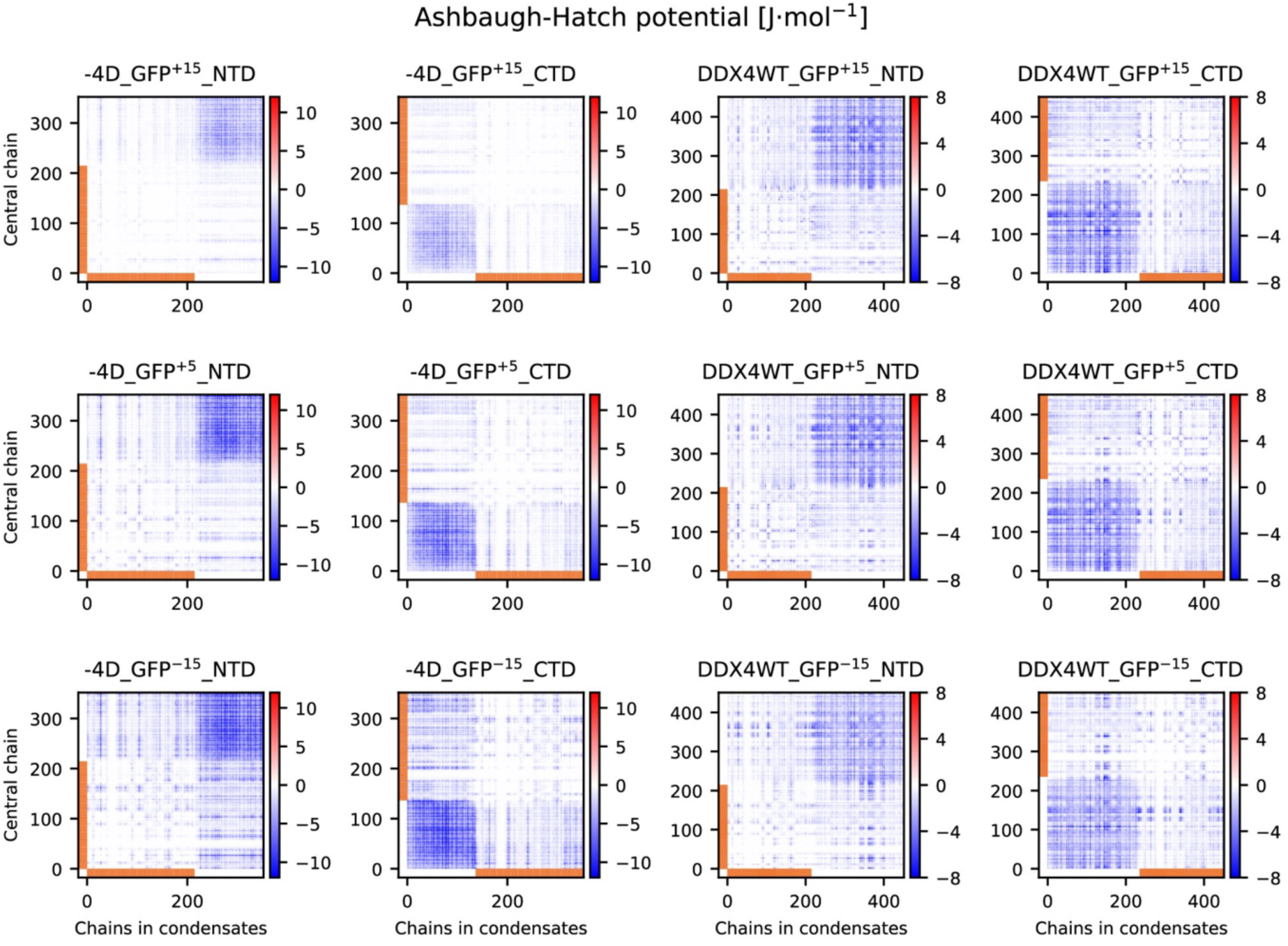
Averaged residue-pair interaction energies (the Ashbaugh-Hatch potential in forcefield) between the most central chain and the rest of the condensate for −4D and DDX4 with three GFP variants. The orange lines indicate folded parts.

**Figure S11:**
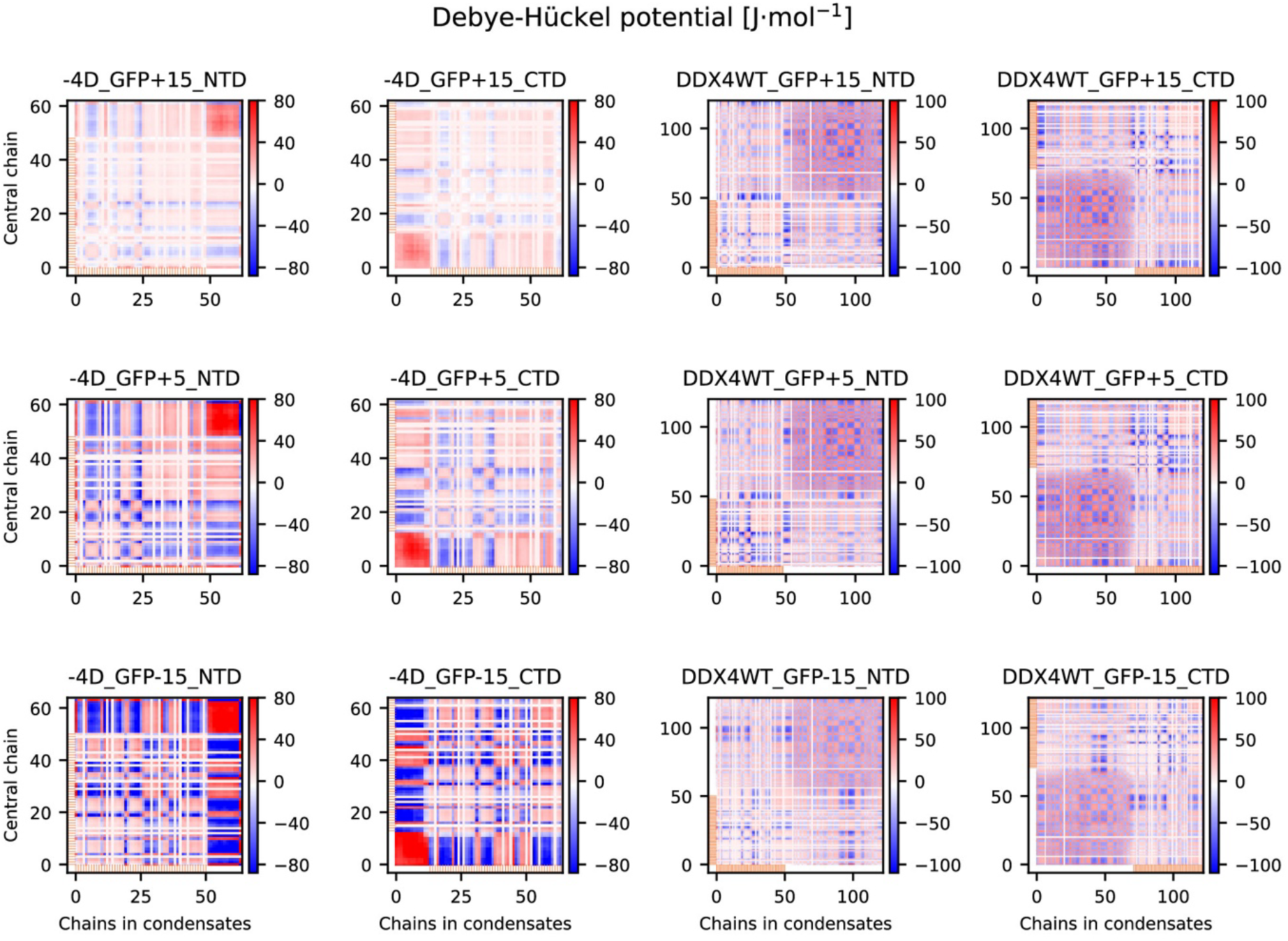
Averaged residue-pair electrostatic interaction energies (the Debye-Hückel potential in forcefield) between the most central chain and the rest of the condensate for −4D and DDX4 with three GFP variants. Only the charged residues are shown. The orange lines indicate folded parts.

## Supplementary tables

**Table S1: Overview of tags used in this study.** Properties of the (fluorescent protein) tags used in this study: including number of amino acids, molecular weight, theoretical pI and extinction coefficients calculated using ExPASy ProtParam; net charge calculated using protpi.ch, monomer/dimer, ExcitationExcitiation/Emission wavelength in nm (Ex/Em), quantum yield (QY), brightness, UniProt ID, GeneBank ID and amino acid sequence. https://web.expasy.org/protparam/, https://www.protpi.ch/Calculator/ProteinTool#Results (advanced using pH 7.5 and ExPASy as source for pKa values), https://www.fpbase.org

**Table S2.** Summary of simulated saturation concentrations (c_sat_) of DDX3X and FUS. Simulations with six different florescent proteins. Protein net charges. NTD and CTD indicate N-terminal and C-terminal tagging respectively. ND: We did not see a stable condensed phase.

**Table S3.** Summary of simulated saturation concentrations (c_sat_) of −4D and DDX4. Simulations with three GFP variants. Protein net charges. NTD and CTD indicate N-terminal and C-terminal tagging respectively. ND: We did not see a stable condensed phase.

**Table S4:** Human cell lines used in this study. Human cell lines used in this study and their source.

**Table S5:** Yeast strains used in this study. Yeast strains used in this study, parental background and genotype.

**Table S6:** Plasmids used in this study. Plasmids and construct details used in this study.

**Table S7:** Protein purification details. For each set of purified proteins, purification steps and storage buffers of purified proteins are listed.

**Table S8:** *In vitro* condensation assay and FRAP assay conditions.

